# Discovery of GluA3 preferring AMPA receptor positive allosteric modulator BRD3290

**DOI:** 10.64898/2026.07.26.740780

**Authors:** Christopher Greaves, William E Martenis, Shawn D Nelson, Jon Madison, Adam Skepner, David Baez-Nieto, Katherine J Stalnaker, Evan P Lebois, Arthur J Campbell, Katie Pelham, Mihaela Magdei, Alex Guletsky, Karen Perez de Arce, Yan-Ling Zhang, Florence F Wagner, Jen Q Pan, Michel Weïwer, Morgan Sheng, Sean P Moran

## Abstract

Schizophrenia is a debilitating neuropsychiatric disease that lacks effective treatments for many symptom domains including negative, cognitive and sleep disturbances. Lack of clear disease etiology has hampered the development of new, effective treatments for the unmet needs of people with schizophrenia. Large scale human genetics have identified rare loss of function mutations that substantially increase risk of developing schizophrenia, including in *GRIA3*, the gene that encodes the GluA3 receptor subunit of the AMPA receptor (AMPAR). Several drug discovery programs have been aimed at developing AMPAR positive allosteric modulators (PAMs) as a novel treatment for schizophrenia. Despite intense drug discovery efforts, there are no FDA approved AMPAR PAMs. We therefore hypothesized that selectively targeting GluA3, the AMPAR subunit implicated by human genetics, could yield a safer and more effective AMPAR PAM for the potential treatment of schizophrenia.

Using a combination of medicinal chemistry, in vitro, and in vivo studies, we discovered BRD3290, a GluA3-preferring AMPAR PAM with reasonable potency in heterologous cells, as well as favorable tolerability and brain exposure. Peripheral administration of BRD3290 engaged an established AMPAR PAM target engagement biomarker but did not improve performance of wildtype mice in the novel object recognition task (NOR), in contrast to the nonselective AMPAR PAM PF-4778574, which improved mouse NOR. These findings suggest that the GluA3 selectivity profile of BRD3290 was insufficient to enhance cognitive function in this mouse NOR paradigm. This work highlights the challenges of AMPAR subtype-selective modulation and provides molecular insights into the ability to develop subtype-selective AMPAR PAMs.

**Graphical Abstract:** 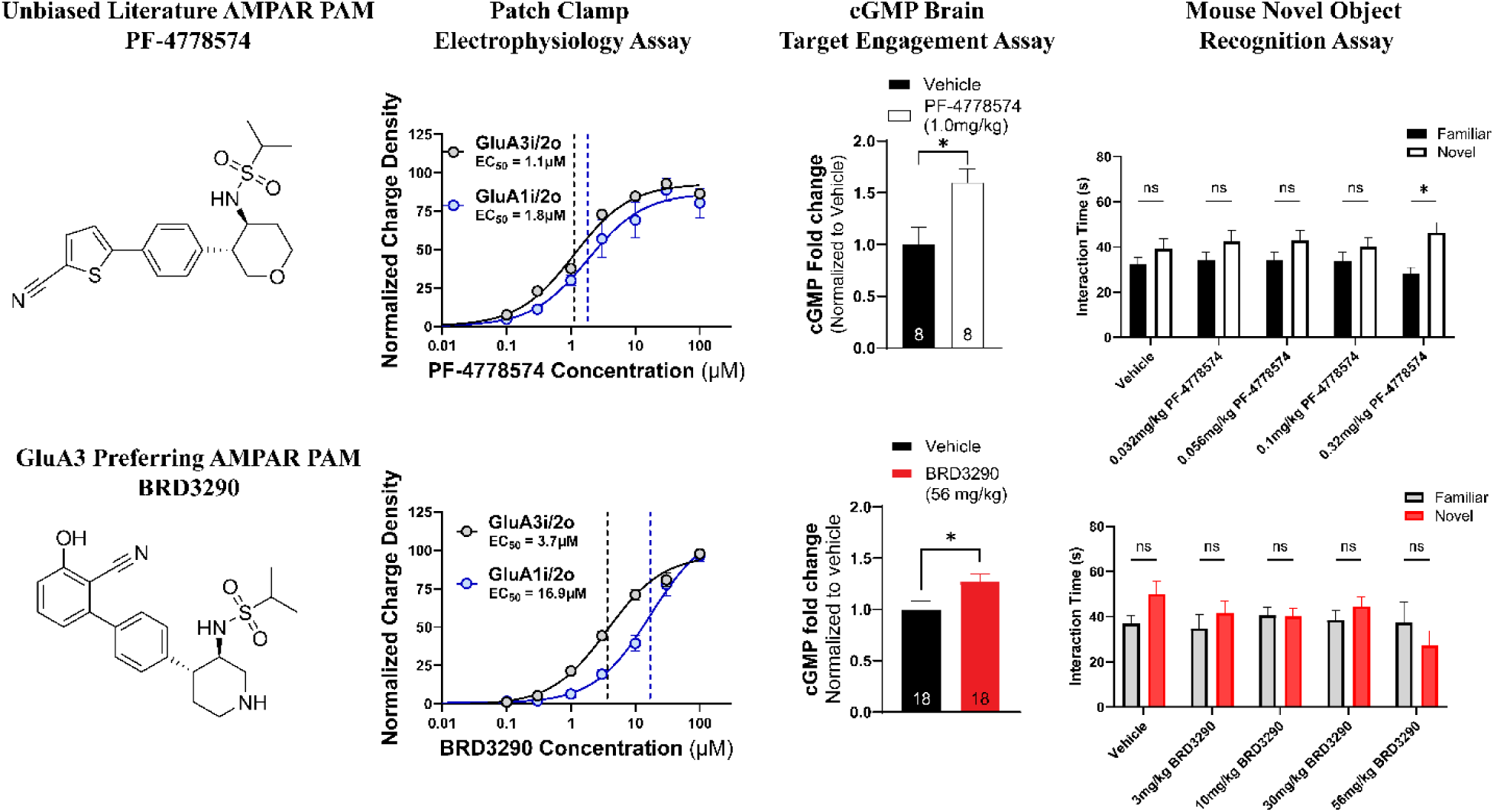

## Introduction

Schizophrenia is a chronic, debilitating psychiatric disorder for which current antipsychotics inadequately address negative symptoms, cognitive impairment, and sleep disturbances. Progress on these unmet needs has been hampered by the disorder’s largely unresolved etiology, which has left drug discovery efforts without clear, mechanistically validated targets. Recent exome sequencing efforts, however, directly associated ultra-rare *GRIA3* loss-of-function variants with increased risk of developing schizophrenia^1^. *GRIA3* encodes the GluA3 (also known as GluR3) protein, an AMPA-type glutamate receptor (AMPAR) subunit^2,3^. AMPARs are postsynaptic ligand-gated ion channels that open upon glutamate binding and are the primary mediators of excitatory synaptic transmission in the brain^3^. AMPARs are obligate tetramers composed of combinations of four different subunit proteins GluA1–4 (encoded by genes *GRIA1*– *GRIA4*), giving rise to functional homotetramers and heterotetramers of which, GluA2-containing heterotetramers are most common in the CNS^2–5^. The rapid activation and deactivation kinetics of AMPARs are critical for shaping excitatory postsynaptic currents, and AMPAR-mediated signaling plays essential roles in synaptic plasticity, learning, and memory^3,6^. Pharmacological enhancement of AMPAR function has therefore been pursued as a strategy to improve cognition in a range of neurological and psychiatric disorders, including Alzheimer’s disease, schizophrenia, depression, attention-deficit disorders^7,8^ and preventing neuroapoptosis ^9^.

Over the past three decades, several structural classes of AMPAR positive allosteric modulators (PAMs) have been discovered^8^. These compounds can modulate AMPAR through various mechanisms including slowing receptor desensitization and/or deactivation^10,11^. Several groups have previously demonstrated that AMPAR PAMs exhibit pro-cognitive efficacy in preclinical animal models^8,12–15^. Despite these encouraging preclinical efficacy results, AMPAR PAMs such as BIIB-104 and PF-4778574 also display serious dose-limiting adverse effects, including convulsions across multiple preclinical species, which result in a narrow therapeutic window^14,15^. This narrow therapeutic window may underlie the lack of efficacy and adverse events seen in the Phase II clinical trial of BIIB-104 in patients with schizophrenia that was recently disclosed by Biogen (ClinicalTrials.gov: NCT03745820). Because *GRIA3* is the only AMPAR gene linked to schizophrenia to date, and GluA3-containing AMPARs represent a smaller subset of total brain AMPARs, we hypothesized that selectively targeting GluA3-containing AMPARs would increase the efficacy and decrease the toxicity of AMPAR PAMs as potential therapeutics for schizophrenia.

Here, we identified BRD3290, a brain-penetrant AMPAR PAM with nearly 10-fold selectivity for GluA3-containing AMPARs over GluA1i/2o in our heterologous cell-based FLIPR assay. Using BRD3290, we set out to directly test our therapeutic hypothesis that a GluA3 selective AMPAR PAM would display a wider therapeutic window than non-subtype selective AMPAR PAMs such as PF-4778574. Although BRD3290 produced no change in the accelerating rotarod fall-latency paradigm when dosed up to 56 mg/kg, a dose that increased brain cGMP (a known AMPAR-dependent biomarker^14–16^), it did not improve novel object recognition in wild-type (WT) mice at any dose tested. This is in stark contrast to PF-4778574, which increased WT mouse NOR at doses slightly below those that produce adverse events. Although BRD3290 did not outperform an established non-selective AMPAR PAM in the NOR assay, it may still prove useful for probing GluA3-selective deficits and behavioral paradigms outside of mouse NOR.

## Results and Discussion

While dozens of AMPAR PAMs have been reported in the literature, little is known about the subtype selectivity of these various AMPAR PAMs. Since our goal was to identify AMPAR PAMs that selectively target GluA3-containing AMPARs, (*GRIA3* is the only AMPAR subunit gene implicated by rare-variant human genetics to date^1^), we first set out to characterize the GluA3 selectivity of previously identified AMPAR PAMs. To accomplish this goal, we developed a set of high-throughput fluorescence cell-based assays to characterize AMPAR activity across eight AMPAR-expressing HEK293 cell lines, each containing different combinations of AMPAR subunits (Figure 1A, B). To streamline the assay and eliminate the usage of fluorescent dyes, each cell line also expressed the genetically encoded calcium indicator GCaMP6s with a modification to enrich expression at the plasma membrane (GCaMP6s-CAAX)^17^.

**Figure 1:**
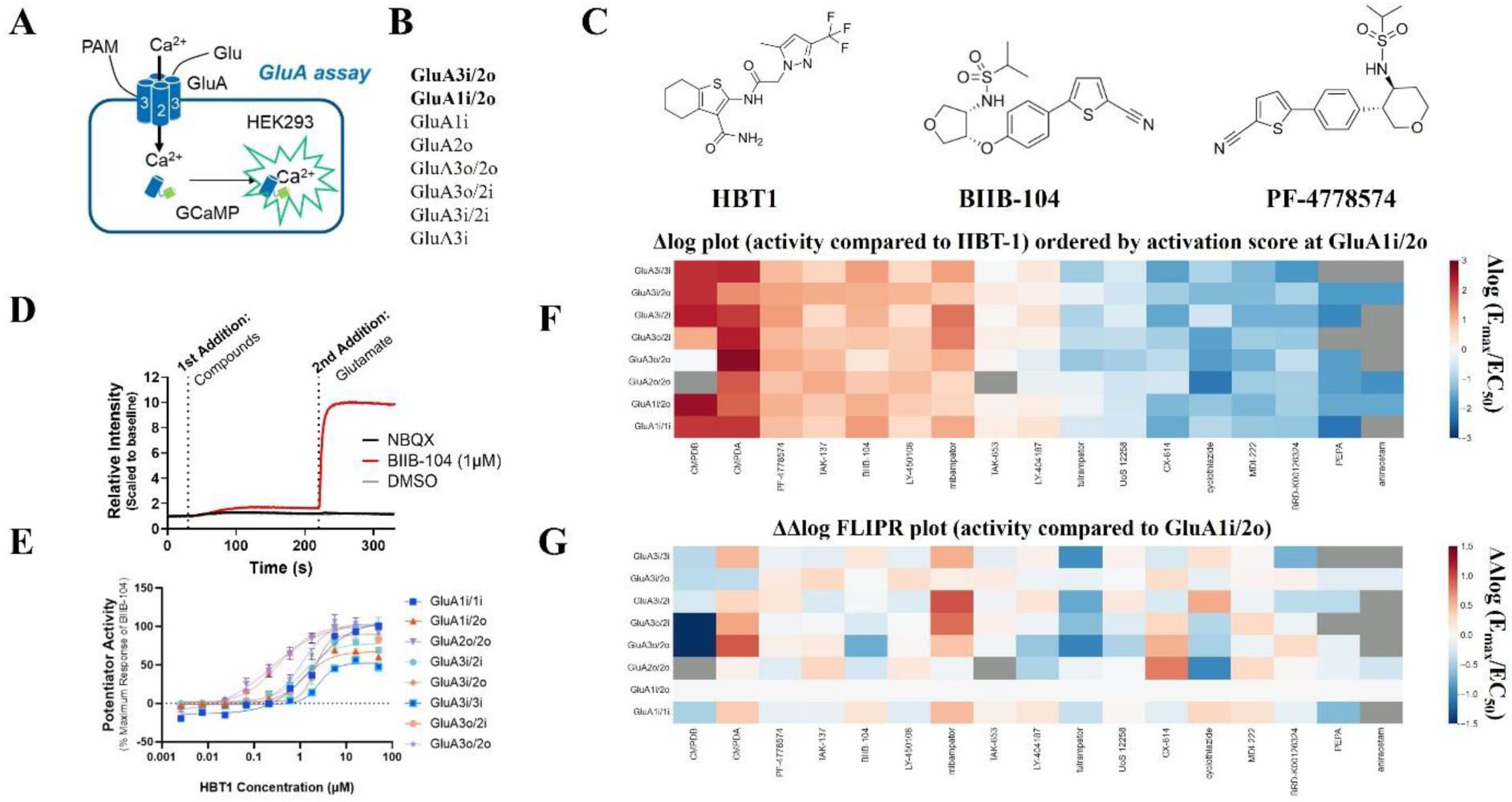
AMPAR cell line assay development and characterization of known potentiators. (A) Fluorescent Imaging Plate Reader (FLIPR®) assay diagram. (B) AMPAR homodimers/heterodimer constructs used to identify selectivity. (C) Molecular structures of key literature comparators. (D) Sample traces of positive, negative, and vehicle controls (BIIB-104, AMPAR inhibitor NBQX, and DMSO, respectively) in FLIPR assay. (E) Sample traces of control compound HBT1 across all eight cell lines in the FLIPR assay. (F) Δlog plot of literature AMPAR potentiators in FLIPR assay (activity in comparison to activity of HBT1 in that cell line). Grey means no activity detected for that compound in that cell line. (G) ΔΔlog plot of literature AMPAR potentiators in FLIPR assay (activity of each compound at a given cell line normalized to the activity of the same compound in the GluA1i/2o cell line). n ≥ 3 biological replicates with each biological replicate consisting of at least 2 same day technical replicates.

AMPARs assemble as obligate tetramers and each subunit is expressed as one of two isoforms, flip (i) and flop (o), each with distinct electrophysiological properties^18^. In our diheteromeric cell lines, consistent stoichiometric assembly was promoted by connecting the two different subunits by a P2A linker (Supplemental Figure 1A). Stable cell line expression of the AMPAR subunits was confirmed via western blot (Supplemental Figure 1B, C). By immunocytochemistry, expression of the AMPAR subunits was negligible in uninduced cell lines, in contrast to the robust plasma membrane expression seen after exposure to doxycycline (Supplemental Figure 1D).

While these eight tetracycline-inducible cell lines do not include all possible combinations of AMPAR subunits, they were chosen based on biological relevance, subunit selectivity, and isoform selectivity. The structures of a subset of literature AMPAR PAMs (HBT1, BIIB-104, and the closely related PF-4778574) are shown in Figure 1C, and a representative FLIPR trace of AMPAR PAM activity in one of our HEK293 FLIPR assays (GluA2i/1o) is shown in Figure 1D. A set of concentration-response curves, for an example AMPAR PAM (HBT1) across all eight cell lines is shown in Figure 1E. Of note, no auxiliary subunits are present in the HEK293 cell lines used in this study; consequently, the fluorescence response to glutamate alone is minimal and cannot be used to set the E_max_ value. To overcome this issue, we instead used the in-plate response of glutamate plus a fixed concentration of the AMPAR PAM BIIB-104 (0.75 µM), which produced a highly consistent response across plates and assays. To create an assay independent value that allowed us to compare selectivity across cell lines^19^, we took the activity score of each compound in the fluorescence assay (E_max_/EC_50_) and compared it against an in-plate reference compound, HBT1 (Δlog (E_max_/EC_50_)) (Figure 1F). HBT1 was selected as our reference compound as it behaved consistently and displayed no intrinsic agonist activity across the various cell lines tested. To understand selectivity across the different cell lines, we used ΔΔlog (E_max_/EC_50_), which normalized activity of a given compound in one cell line to the activity of that same compound in the non-GluA3 containing AMPAR GluA1i/2o cell line (Figure 1G).

The cell line expressing GluA1i/2o was chosen as the “anti-target” since AMPARs containing GluA1 and GluA2 subunits comprise the majority of GluA3-lacking AMPARs in brain regions important for learning and memory^5^. The cell line expressing GluA3i/2o was chosen as the target cell line for several reasons. Firstly, GluA3i mRNA expression is higher than GluA3o expression in brain regions critical for cognition such as the cortex, hippocampus and basal ganglia based on GTEx brain expression data ([GTEx Portal](https://gtexportal.org/home/), dbGaP accession number phs000424.vN.pN). Secondly, there is limited evidence that GluA3 homomers are expressed in native brain tissue^20^. Thirdly, most AMPARs contain at least one GluA2 subunit and previous studies have shown that GluA3 subunits are predominantly associated with GluA2 in brain tissue^2,4,5,11^.

These results (Figure 1G) demonstrate that there are different selectivity patterns across the different AMPAR PAMs tested; however, no previously disclosed AMPAR PAM displayed substantial selectivity for the GluA3-containing AMPAR cell line GluA3i/2o in the fluorescence assay. In the absence of any existing compounds with good GluA3i/2o selectivity, we chose PF-4778574 as the starting point for our own medicinal chemistry campaign, with the aim of introducing the desired selectivity while retaining the favorable potency and brain penetrance of this scaffold^15^.

At 0.068 µM, PF-4778574 displayed similar activity in both the GluA3i/2o (Figure 2A) and GluA1i/2o (Figure 2B) FLIPR assays. This is further illustrated by the negligible selectivity window between GluA3i/2o and GluA1i/2o for PF-4778574 (Figure 2C). We explored different replacements for the cyanothiophene moiety on both tetrahydropyran and piperidine cores. (Table 1). Movement of the heteroatom in the saturated ring led to only moderate losses in activity in the FLIPR assays (compound **7**). 4-hydroxy, methoxy or cyanophenyl substituents all afforded nonselective compounds with low overall potencies. Positioning a cyano group *ortho* to the aryl-aryl linkage (**5**, **6**, **11**, **12**) afforded relatively more potent compounds where some selectivity over GluA1i/2o and GluA1i/1i emerged, hinting that this position might provide a way to improve GluA3 selectivity.

**Figure 2:**
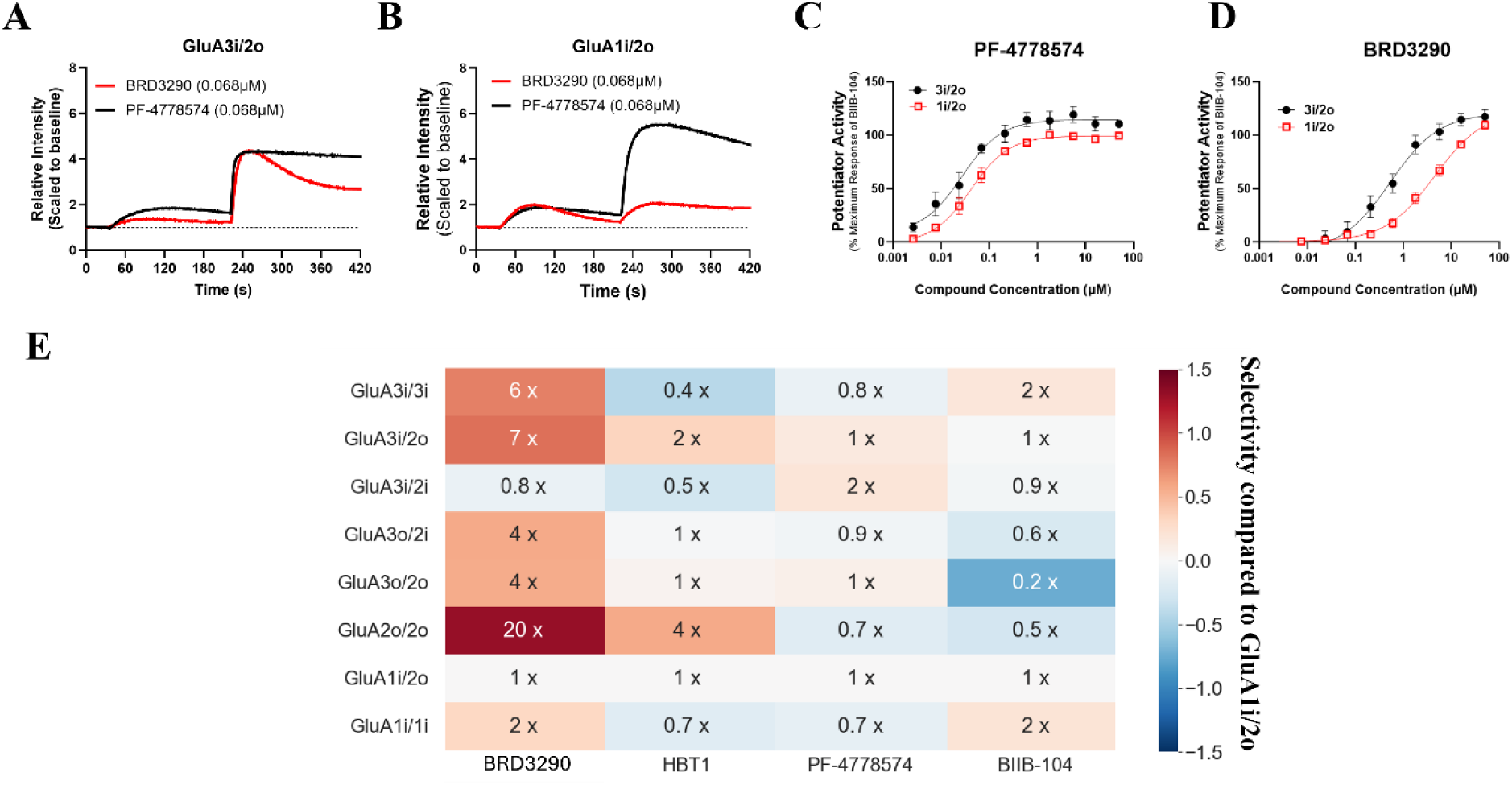
High throughput fluorescence in vitro cell-based assay to identify GluA3-preferring AMPAR PAMs. Representative traces of BRD3290 (red) and PF-4778574 (black) response in the (A) GluA3i/2o and (B) GluA1i/2o FLIPR cell lines. Averaged responses to (C) PF-4778574 and (D) BRD3290 in 1i/2o (black) and 3i/2o (red) cell lines in FLIPR assay. (E) ΔΔlog activity plot of BRD3290 and select literature AMPAR potentiators in FLIPR assay (activity of each compound at a given cell line normalized to the activity of the same compound in the GluA1i/2o cell line). Data points display mean ± SEM of n ≥ 3 biological replicates with each biological replicate consisting of at least 2 same day technical replicates.

**Table 1:** Activity and selectivity data for SAR. Max selectivity is the highest fold selectivity observed for any GluA3-containing heterodimer compared to GluA1i/2o. Average Δlog is the arithmetic mean of all single Δlog unit activity measurements referenced to HBT-1 across all cell lines measured (a Δlog value of 1.0 indicates a compound that is 10 x more active than HBT-1). Full details are in Supplemental Tables S1 and S2.

|  |  | R-group |  |  |  |  |  |
| --- | --- | --- | --- | --- | --- | --- | --- |
| Piperidine | Compound no. | **1** | **2** | **3** | **4** | **5** | **6** |
|  | Max selectivity for any GluA3 containing cell line | 1 x | 1 x | 2 x | 0.9 x | 2 x | 3 x |
| | Average $\Delta\log$ activity vs. HBT-1 | -0.52 | -0.71 | -0.33 | -0.79 | -0.87 | 0.96 |

**Table 1:** Activity and selectivity data for SAR.
| THP | Compound no. | 7 | 8 | 9 | 10 | 11 | 12 |
| --- | --- | --- | --- | --- | --- | --- | --- |
|  | Max selectivity for any GluA3 containing cell line | 1 x | 2 x | 3 x | 0.8 x | 5 x | 2 x |
| | Average $\Delta\log$ activity vs. HBT-1 | 0.75 | 0.46 | -0.21 | 0.02 | 0.26 | 1.77 |

We further examined substituents at the 2- and 3-positions relative to the aryl linkage in the piperidine series (Table 2). Benzoxazole **13** afforded a weaker compound which retained some GluA3 selectivity. An attempt to prepare the matched benzisoxazole instead afforded the ortho-hydroxybenzonitrile BRD3290, a consequence of rupture of the N-O bond during the cross coupling. This compound demonstrated substantially higher activity in the GluA3i/2o FLIPR assay (Figure 2A) than in the GluA1i/2o FLIPR assay (Figure 2B), representing a larger selectivity window (Figure 2D) for a GluA3-containing cell line over a GluA3-lacking cell line. Based on fold change in activity over the GluA1i/2o cell line, we demonstrated the improved GluA3 selectivity of BRD3290 compared to a set of previously reported compounds (Figure 2E).

**Table 2:**
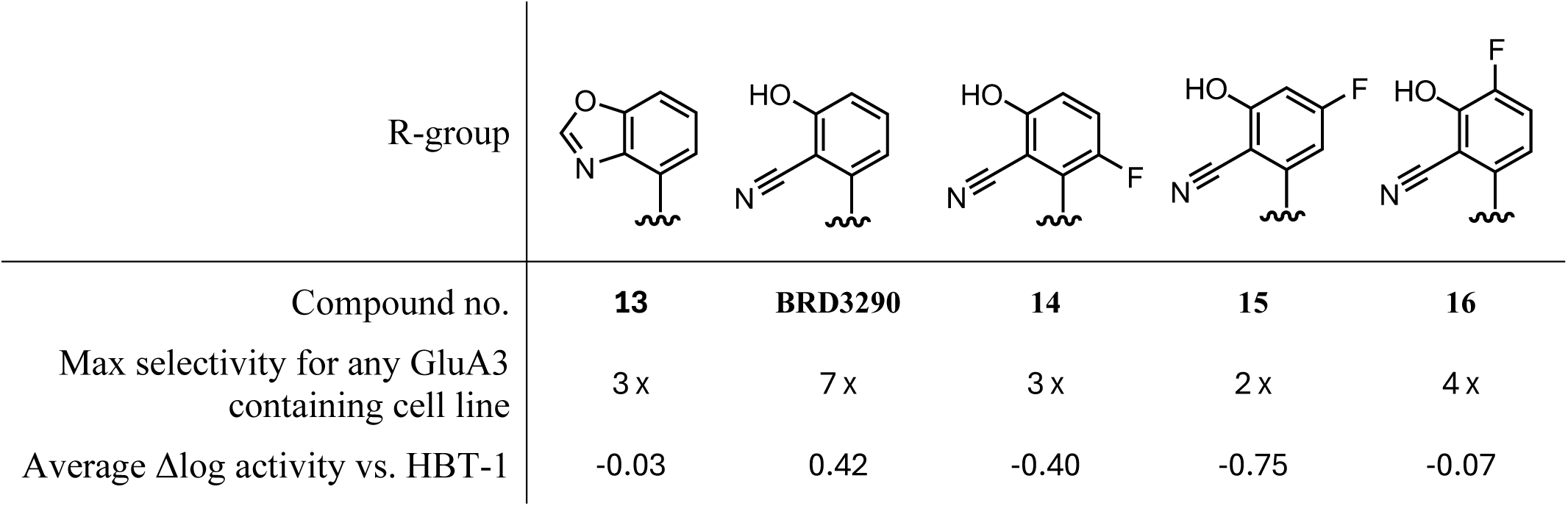
Additional activity data on the piperidine series.

| R-group |  |  |  |  |  |
| --- | --- | --- | --- | --- | --- |
| Compound no. | **13** | **BRD3290** | **14** | **15** | **16** |
| Max selectivity for any GluA3 containing cell line | 3 x | 7 x | 3 x | 2 x | 4 x |
| Average Δlog activity vs. HBT-1 | -0.03 | 0.42 | -0.40 | -0.75 | -0.07 |

Modest changes to BRD3290 decreased selectivity and potency, as in the fluorinated series **14**, **15**, and **16**, and analogues in Supplemental Table S3. BRD3290 remained the most GluA3 selective compound in our series, so we explored alternative synthetic routes to generate sufficient material for secondary assays.

Reported synthetic procedures^21^ for this scaffold are lengthy and require many steps to establish the correct stereochemistry on the piperidine ring. We sought a more efficient route to enable preparation of larger quantities of material for in vivo studies. We speculated that reaction of a sulfonylated aziridine with an aryl Grignard reagent would allow the installation of the 4-aryl group while revealing the desired sulfonamide moiety, whose electron-withdrawing character would also facilitate the ring opening. Towards this end, sulfonyl aziridine **18** was prepared from racemic piperidine epoxide **17**.

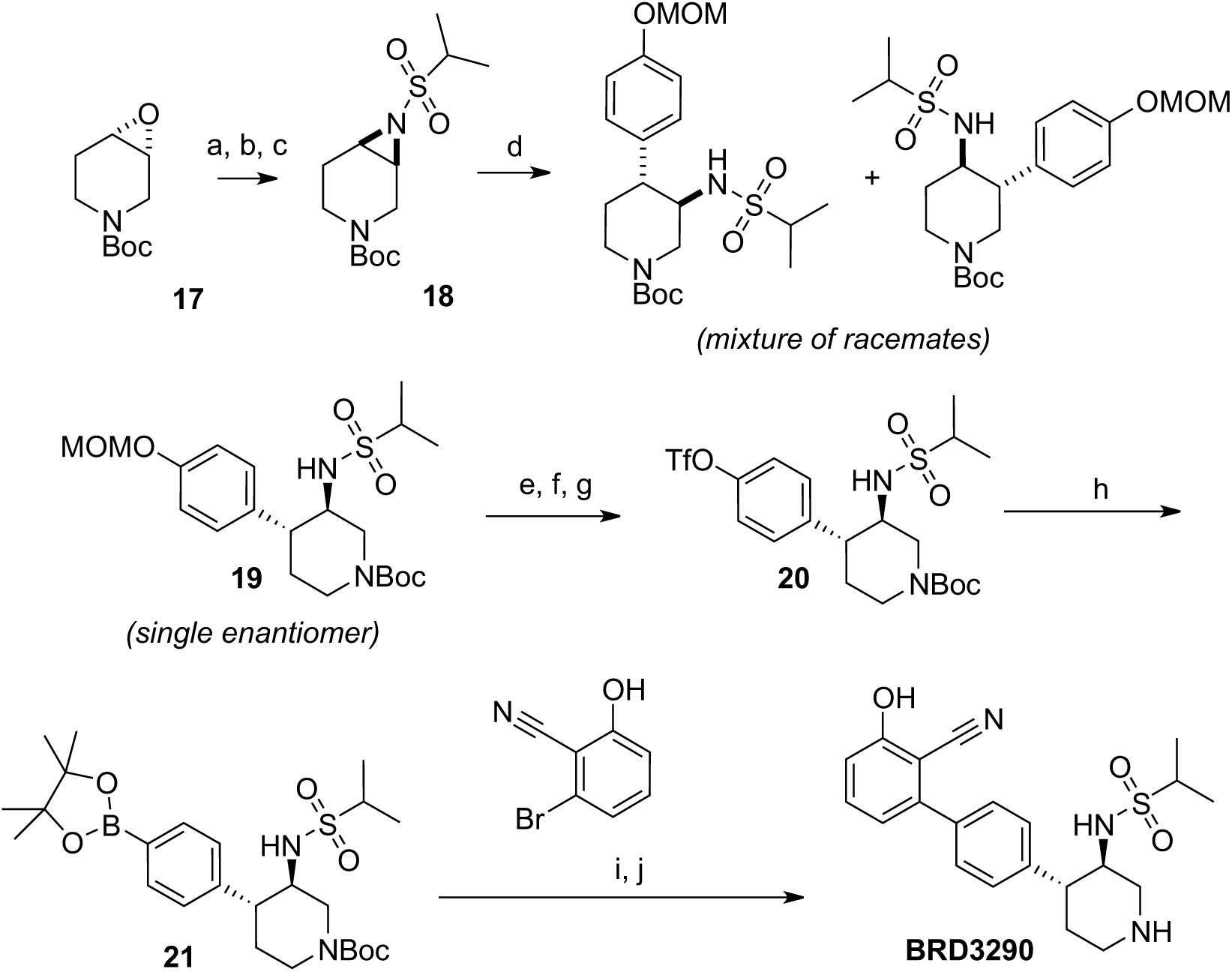

Reagents and conditions: (a) iPrSO_2_NH_2_, K_2_CO_3_, TEBAC, dioxane (b) MsCl, pyridine, DCM (c) K_2_CO_3_, MeCN; (d) ArMgBr, Cu(OTf)_2_, THF; (e) HCl, dioxane (f) Boc_2_O, Et_3_N, MeOH (g) Tf_2_O, pyridine, DCM; (h) bis-pin, KOAc, Pd(dppf)Cl_2_, dioxane; (i) ArBr, K_3_PO_4_, Pd(PPh_3_)_4_, dioxane, H_2_O (j) HCl, dioxane.

Treatment of **18** with the appropriate methoxymethyl-protected Grignard reagent in the presence of copper triflate afforded a mixture of two diastereomeric racemates, a reaction which could be carried out on a 10-gram scale, and which preferentially afforded the desired regioisomer in a roughly 4:1 ratio. Silica chromatography followed by supercritical fluid chromatography over a chiral stationary phase with CO_2_-MeOH as the eluent afforded single enantiomer **19** in 39% overall yield from the starting aziridine. Deprotection, conversion to the boronic ester **21** by way of triflate **20**, and subsequent cross-coupling, afforded BRD3290. Using this route, 10 grams of BRD3290 were prepared, enabling extensive characterization of this compound in secondary in vitro and in vivo assays.

Changes in fluorescence measured in the FLIPR cell-based assay are an indirect measurement of AMPAR channel activity. We therefore performed a secondary assay to directly measure AMPAR PAM changes to channel activity utilizing a 384-well format automated patch clamp electrophysiology assay in each of the eight AMPAR cell lines. Using a ligand-puff addition protocol (Figure 3A), we could rapidly add and remove ligands from the patched cells, thereby mimicking a large glutamatergic synaptic input. Glutamate addition produced a concentration dependent increase in current in each cell line, represented by the GluA3i/2o expressing HEK293 cell line response shown in Figure 3B. Since these cell lines lack the AMPAR auxiliary proteins, we observed a modest response to glutamate similar to what is observed in the FLIPR assay. In contrast to glutamate alone, glutamate in the presence of an AMPAR PAM such as BIIB-104 generated a large concentration-dependent increase in AMPAR currents (Figure 3C). HBT1 (Figure 3D), BIIB-104 (Figure 3E) and PF-4778574 (Figure 3F) showed relatively minimal differences between EC_50_ values across the GluA3-containing cell line GluA3i/2o compared to the GluA3-lacking cell line GluA1i/2o. BRD3290 displayed a substantially larger selectivity window between the EC_50_ of the two key cell lines (Figure 3G), reproducing the increased GluA3 selectivity observed in the FLIPR-based assay. Selectivity across all eight cell lines is shown in Table 3. BRD3290 demonstrated relatively similar selectivity windows for GluA3i/2o over GluA1i/2o in our electrophysiology assay compared to the FLIPR based assay (Figure 2E).

**Figure 3:**
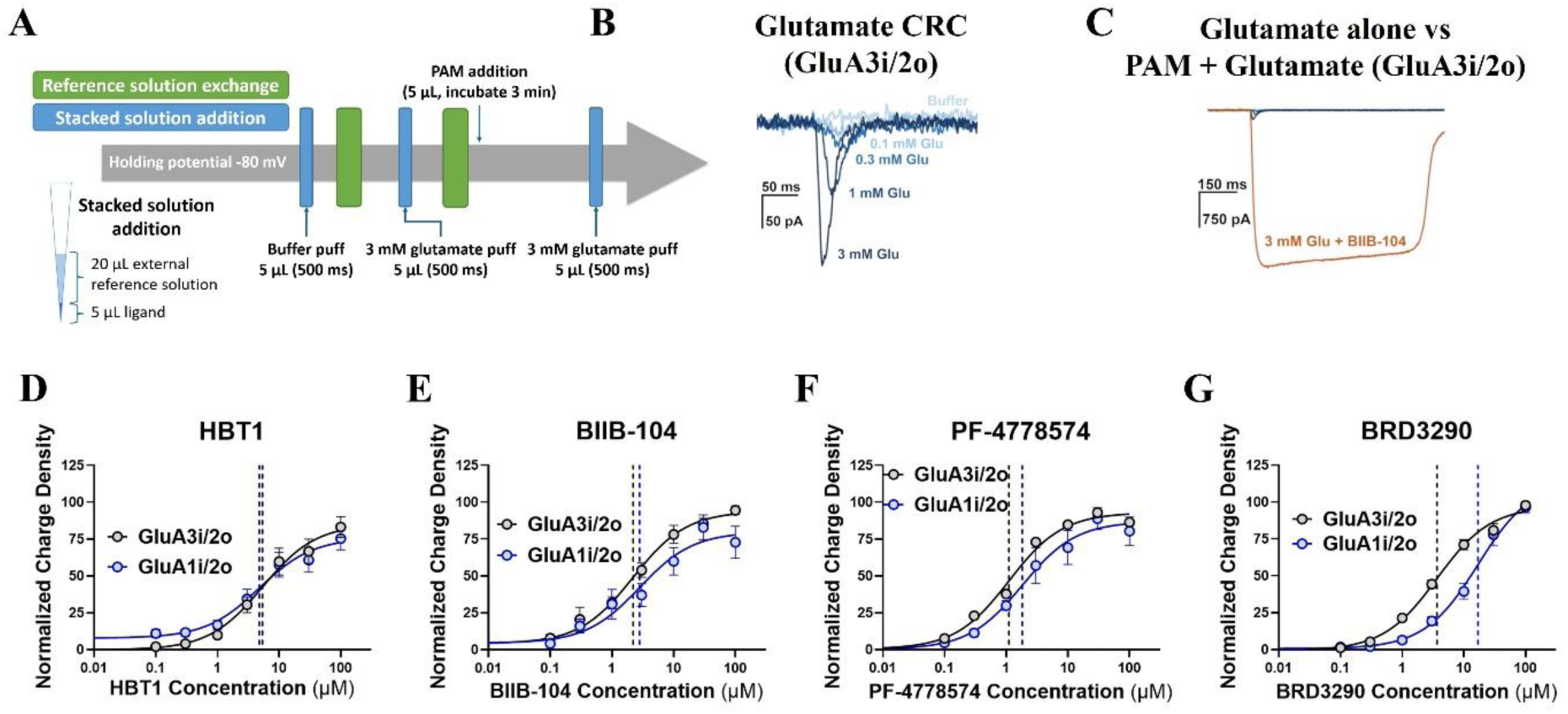
Characterization of AMPAR PAMs using automated patch clamp electrophysiology assay. (A) Automated patch clam electrophysiology assay diagram. Stacked ligand (glutamate or extracellular solution) delivered over 500 ms. AMPAR PAMs were added in concentration-response curve across 6 half-log steps. (B) Representative concentration-response curves for 3 mM glutamate alone and (C) 3 mM glutamate + 1 µM BIIB-104 in the GluA1i/2o cell line. (D-G), Concentration-response curves detailing charge density response normalized to the average peak charge density per replicate for each compound at each cell line. Vertical dashed lines denote EC_50_ for each compound in each cell line; all values in Table 3. Data are displayed as mean ± SEM consisting of n ≥ 3 biological replicates, with each biological replicate consisting of at least 3 cells for each compound per concentration across all 8 cell lines.

**Table 3:**
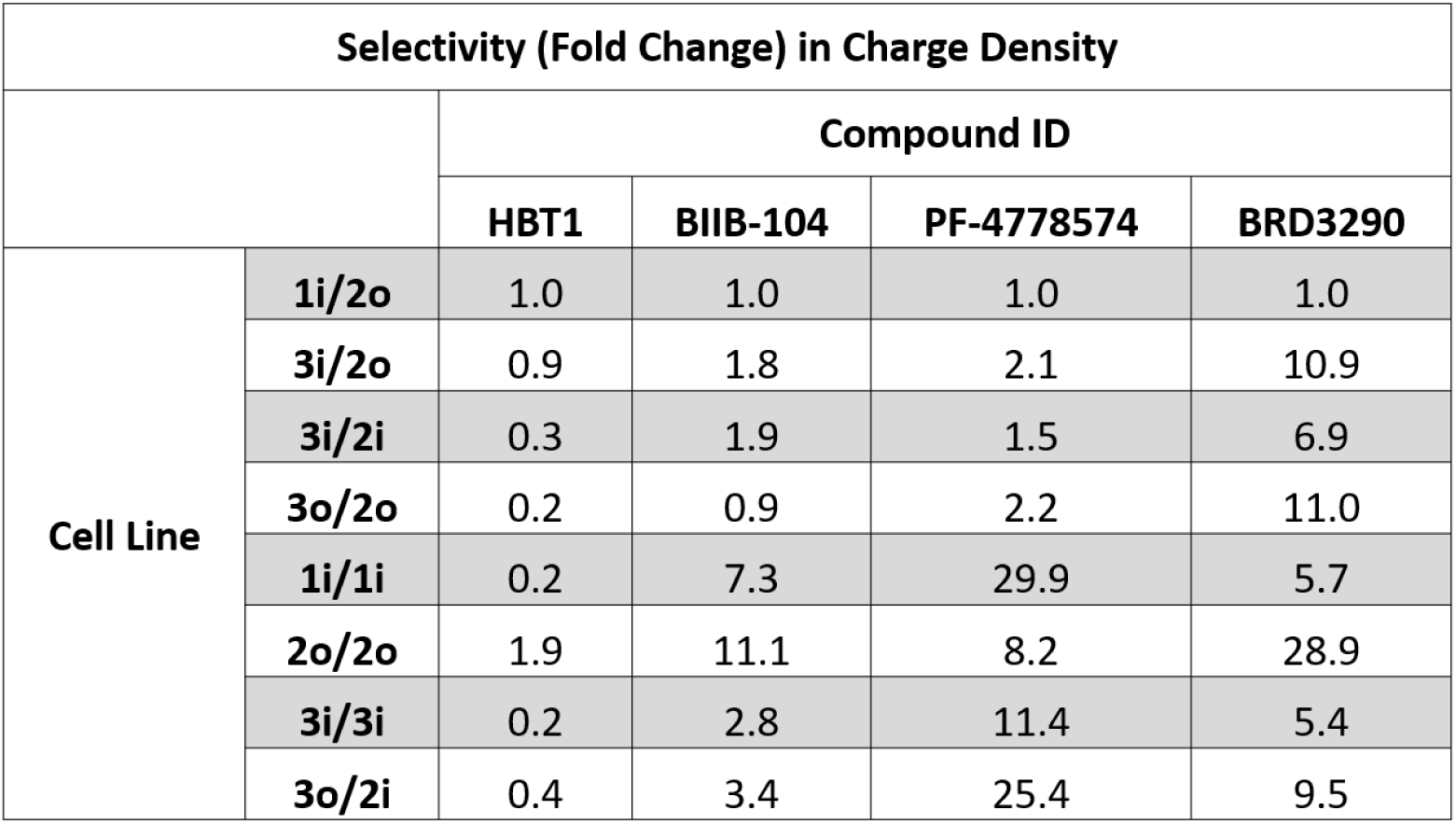
Automated Patch Clamp Electrophysiology results of key AMPAR PAMs.

| Selectivity (Fold Change) in Charge Density |  |  |  |  |  |
| --- | --- | --- | --- | --- | --- |
|  |  | Compound ID |  |  |  |
|  |  | HBT1 | BIIB-104 | PF-4778574 | BRD3290 |
| Cell Line | 1i/2o | 1.0 | 1.0 | 1.0 | 1.0 |
|  | 3i/2o | 0.9 | 1.8 | 2.1 | 10.9 |
|  | 3i/2i | 0.3 | 1.9 | 1.5 | 6.9 |
|  | 3o/2o | 0.2 | 0.9 | 2.2 | 11.0 |
|  | 1i/1i | 0.2 | 7.3 | 29.9 | 5.7 |
|  | 2o/2o | 1.9 | 11.1 | 8.2 | 28.9 |
|  | 3i/3i | 0.2 | 2.8 | 11.4 | 5.4 |
|  | 3o/2i | 0.4 | 3.4 | 25.4 | 9.5 |

We next confirmed that increasing the GluA3 selectivity of BRD3290 did not substantially alter the favorable drug-like properties of the parent compound PF-4778574, namely high aqueous solubility, low clearance, and high brain exposure. BRD3290 retained tolerable in vitro mouse microsome clearance, high aqueous solubility and a reasonable P-glycoprotein (PGP)-efflux ratio (Supplemental Table S4). These favorable in vitro DMPK values for BRD3290 translated into relatively high and sustained unbound mouse brain exposures that were suitable to facilitate subsequent in vivo testing (Figure 4A, Supplemental Figure 2).

**Figure 4:**
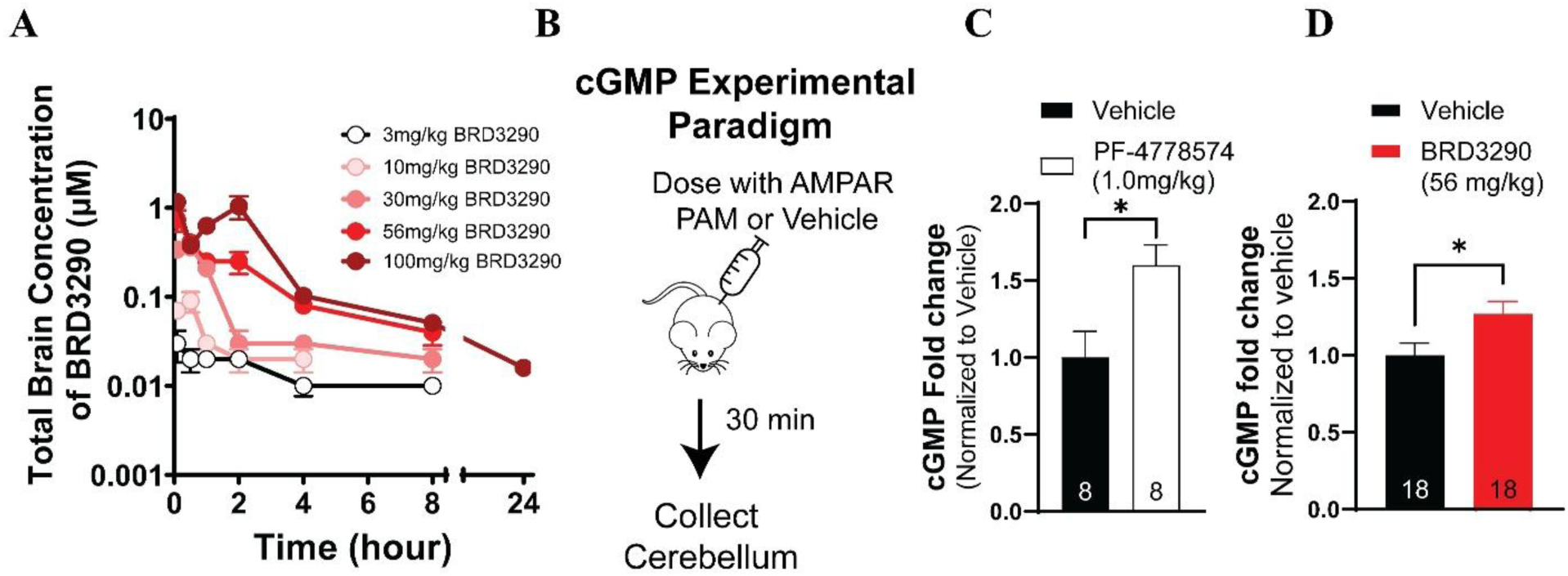
BRD3290 is a brain penetrant GluA3 AMPAR PAM that increases cerebellar cGMP. (A) Total unbound BDR3290 in brain lysate as quantified by LC-MS/MS following IP injection, n = 3 for each data point method (lower limit of quantification 2.04 ng/mL; data points below LLOQ not displayed for clarity, though data collected for all doses at all timepoints). (B) Visual representation of cGMP assay for target engagement. Quantification of cGMP in mouse cerebral cortex by ELISA upon highest dose tested (C) PF-4778574 or (D) BRD3290, with samples normalized for cerebellum weight and measured in technical triplicate. Data displayed as mean ± SEM, *p<0.05. Student’s t-test. Number of animals per group displayed in each bar.

It was previously shown that AMPAR receptor PAMs such as BIIB-104 and PF-4778574 can induce severe adverse events such as convulsions in preclinical species^14,15^. Therefore, we set out to compare the adverse event liability of our GluA3-preferring AMPAR PAM BRD3290 to PF-4778574 utilizing the modified Racine score^22^. PF-4778574 at 1.78 mg/kg, but not 1.0 mg/kg, produced convulsions, in complete agreement with previously published results (Table 2)^15^. We then set out to test BRD3290 in the behavioral convulsion assay across a wide range of doses guided by our PK studies. BRD3290 did not result in convulsions at doses tested up to 100 mg/kg in mice (Table 2). Since administration of 100 mg/kg BRD3290 did not produce significantly higher C_b,u_ or cerebrospinal fluid (CSF) levels than 56 mg/kg, and neither dose resulted in convulsions, we limited our highest dose of BRD3290 to 56 mg/kg. The lack of seizure liability seen with high doses of BRD3290 could be explained by either a larger safety window or a lack of target engagement.

**Table 2:**
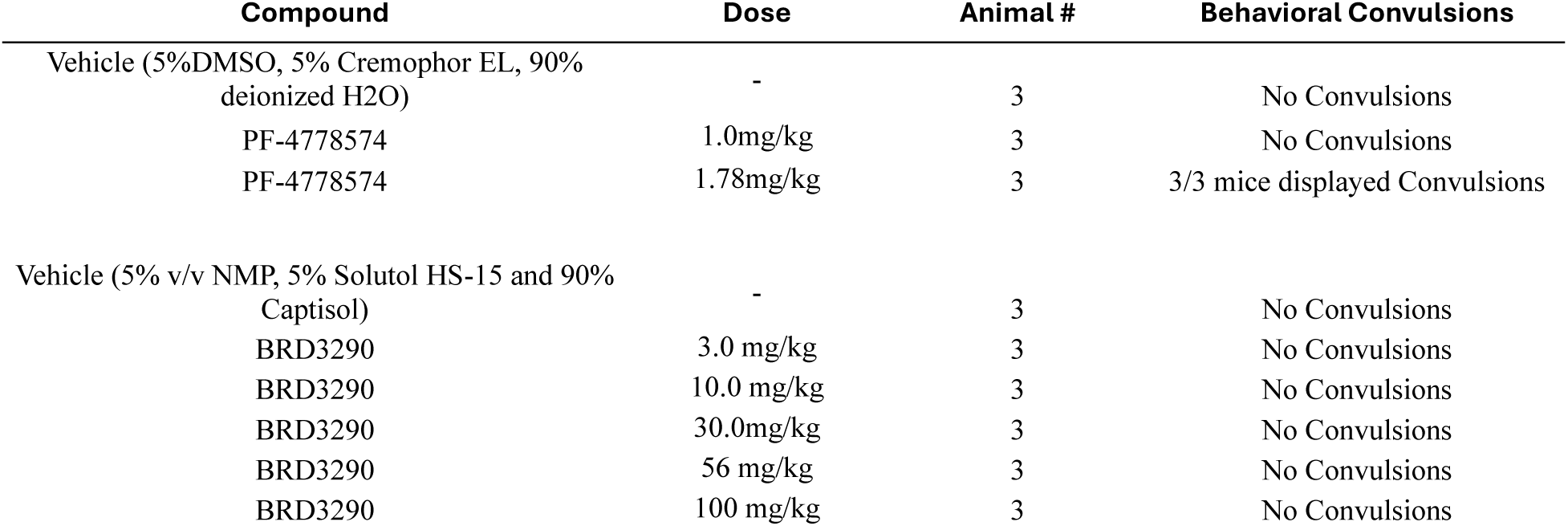
Convulsion liability of PF-4778574 and BRD3290 in mice.

Since previous reports have demonstrated that peripheral administration of AMPAR PAMs can increase cGMP levels in mouse cerebellum^14–16^, we tested the ability of both PF-4778574 and BRD3290 to modulate this AMPAR-dependent target engagement biomarker. Similar to the previously published results^15^, the non-subunit selective AMPAR PAM PF-4778574 resulted in an increase in mouse cerebellar cGMP when dosed at 1.0 mg/kg (Figure 4B, C). BRD3290 also increased mouse cerebellar cGMP after administration of 56 mg/kg (Figure 4D). These results demonstrate that the GluA3-preferring AMPAR PAM BRD3290 can increase an AMPAR-dependent target engagement biomarker in a similar fashion to the non-subunit selective AMPAR PAM PF-4778574, albeit at different doses.

Importantly, previously reported AMPAR PAM adverse events are not limited to convulsions, and changes in motor function have been demonstrated at doses below those that induce overt convulsions^14,15^. Therefore, we compared the effects of PF-4778574 and BRD3290 on gross motor function by utilizing the accelerating rotarod assay (Figure 5A). Decreases in rotarod fall latency can indicate changes to motor function, arousal, or both. PF-4778574 at 0.56 mg/kg, a dose below that required to induce overt convulsions, produced a statistically significant decrease in rotarod fall latency relative to vehicle-treated mice (Figure 5B). Importantly, BRD3290 did not significantly alter fall latency, relative to vehicle-treated mice, at the same dose (56 mg/kg) that significantly changed the AMPAR-dependent biomarker (Figure 5C, Figure 4D).

**Figure 5:**
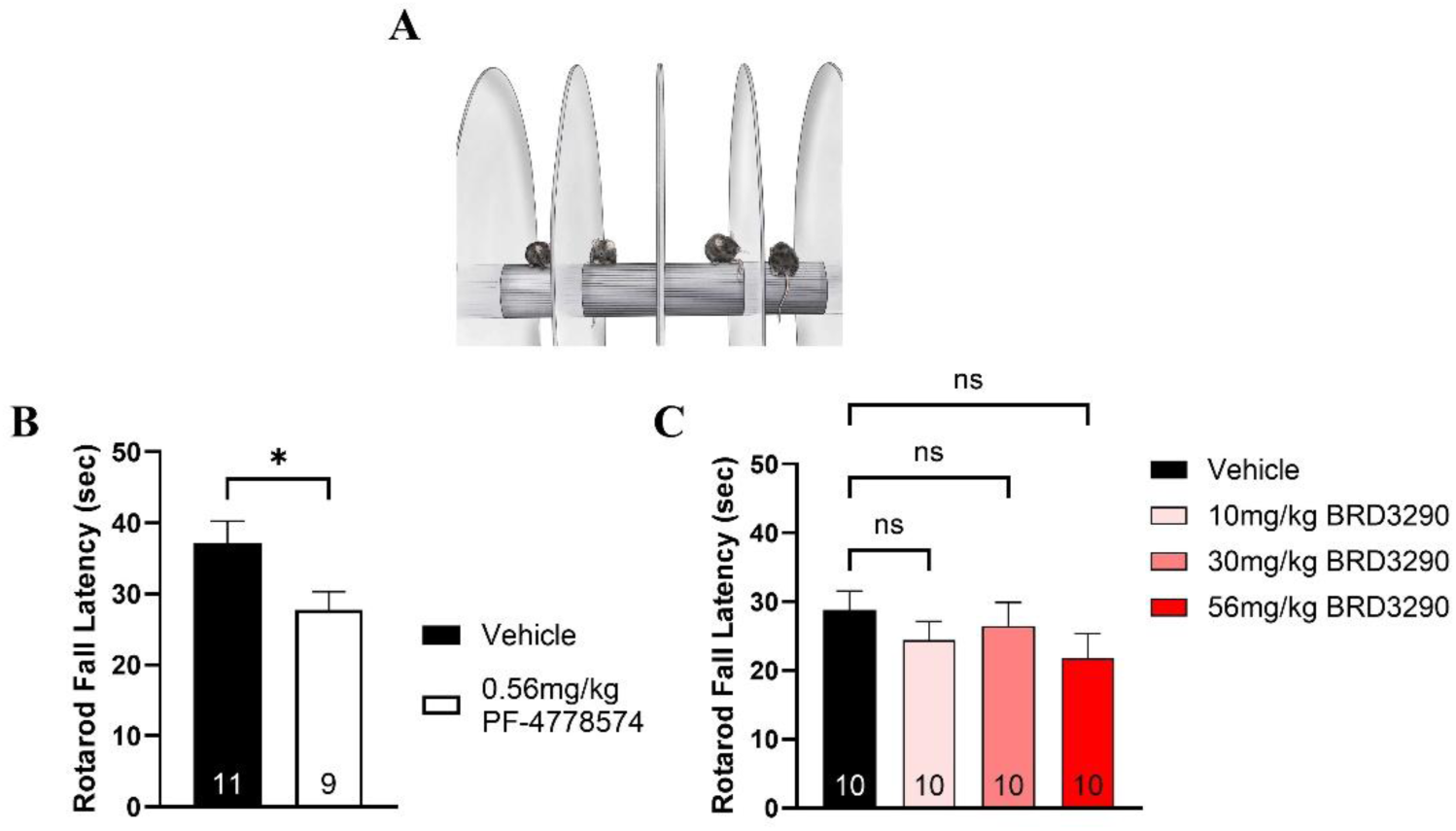
Adverse event liability of PF-4778574 and BRD3290 in WT mice. (A) Visual representation of rotarod assay. (B) AMPAR PAM PF-4778574 at 0.56 mg/kg resulted in a decreased fall latency in mice. (C) BRD3290 does not cause motor deficits at or below 56 mg/kg. Data displayed as mean ± SEM, *p<0.05. Student’s t-test or one-way ANOVA with Dunnett’s post hoc test. Number of animals per group displayed in each bar.

We then compared whether the GluA3-preferring AMPAR PAM BRD3290 performed better than PF-4778574 in a well-characterized rodent cognition assay that is sensitive to AMPAR PAMs. Previous studies have shown that an AMPAR PAM can rescue MK-801 induced deficits in spontaneous alternation^13^. Unfortunately, we could not detect a rescue of MK-801 induced deficits using any dose of the AMPAR PAM PF-4778574 (Supplemental Figure 3A-C). Since several AMPAR PAMs have shown efficacy in increasing novel object recognition in rodents^12,23^, a rodent model of object memory, we then set out to compare the effects of PF-4778574 and BRD3290 in the WT mouse novel object recognition paradigm. Naïve mice were dosed with either vehicle or AMPAR PAM and then allowed to freely explore two identical objects. 24 hr later, mice were returned to the same box, but one of the familiar objects was replaced with a novel object (Figure 6A). PF-4778574 at 0.32 mg/kg, but not at lower doses, produced a statistically significant increase in time spent exploring the novel versus the familiar object (Figure 6B). We did not test higher doses of PF-4778574 since 0.56 mg/kg PF-4778574 produces rotarod fall-latency changes that would confound this rodent cognition assay. In contrast to PF-4778574, no dose of BRD3290 tested resulted in an increase in novel object recognition (Figure 6C). These data demonstrate that although 56 mg/kg BRD3290 increased the AMPAR-dependent target engagement biomarker, it did not improve performance in this 24 hr rodent object recognition memory task.

**Figure 6:**
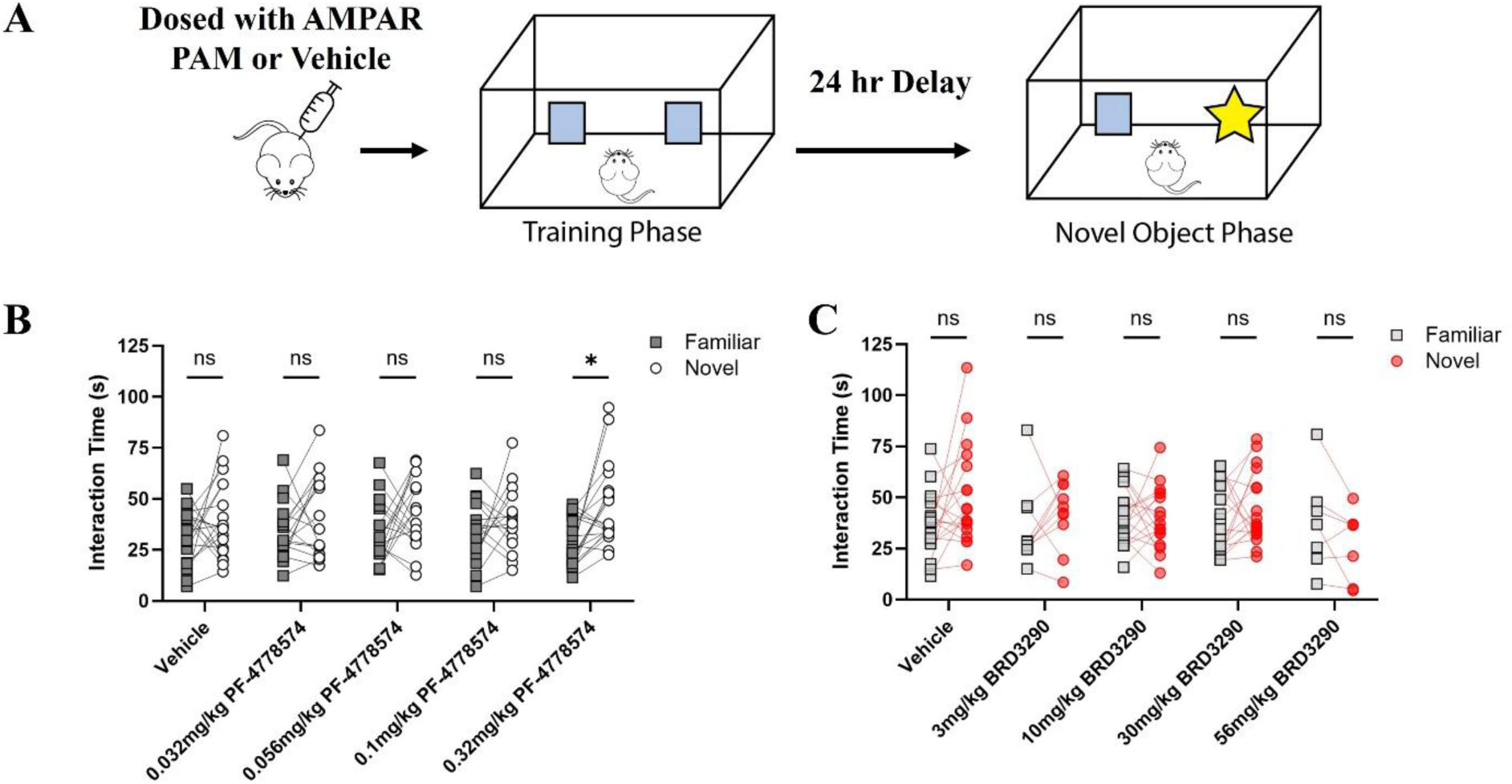
Effects of distinct AMPAR PAMs on 24 hr novel object recognition in WT mice. (A) Visual representation of novel object interaction assay. (B) Time spent exploring the novel (white circle) or the familiar object (grey box) for vehicle treated or PF-4778574 treated mice. (C) Time spent exploring the novel (red circle) or the familiar object (grey box) for vehicle treated or BRD3290 treated mice. Multiple t-tests with multiple comparison correction. *p<0.05.

## Conclusion

BRD3290 is a GluA3 subunit-preferring AMPAR PAM, the first in vitro subunit-selective AMPAR PAM reported in the literature. It is brain-penetrant and engages cerebellar AMPARs at a dose below its toxicity threshold, but no cognitive benefit was seen, at least in the mouse NOR paradigm. Although these data do not support the hypothesis that this GluA3-preferring AMPAR PAM with modest selectivity (∼10-fold GluA3i/2o over GluA1i/2o) would provide benefit in the cognitive domains of schizophrenia as assessed by the rodent NOR assay, subunit-selective AMPAR PAMs could still ameliorate other aspects of the disease not captured by rodent NOR, such as performance in rule-based learning tasks or in aggression assays^24^. These data also do not rule out that molecules with different GluA3 selectivity profiles (e.g., selectively potentiating GluA3 homomers) or larger selectivity windows (e.g. 100-fold fold GluA3i/2o over GluA1i/2o) could prove effective at improving cognition while minimizing on-target toxicity. It also remains to be determined whether GluA3-preferring molecules such as BRD3290 could be beneficial in disorders driven by GluA3 hypofunction^25–27^. Notably, this work provides comprehensive evidence that substantial subunit selectivity is absent across all literature AMPAR PAMs tested but that development of subunit selective compounds is a real possibility and that future studies may be able to extend this subunit selectivity even further. Lastly, these data demonstrate BRD3290’s utility as a tool compound to study GluA3-containing AMPARs in vitro and in vivo as well as serving as a proof of concept for the idea that some degree of subunit selectivity can be engineered into these molecules without sacrificing other drug-like parameters such as potency and brain exposure.

## Materials and Methods

### Study design

Sample size was determined by a priori power analysis, while minimizing the number of animals based on previous experience in each behavioral assay. Sample numbers for all studies can be found in either the figure itself or in the figure legends. All animal studies were performed under approval of the Broad Institute of MIT and Harvard Institutional Animal Care and Use Committee (IACUC). No assessment, or adjustment of, outliers took place in any dataset and all data are displayed. Investigators were blinded to compound treatment for cell-based assays such as FLIPR and for animal behavior studies during both the data collection and data analysis phases.

### Brain Pharmacokinetic Studies

Brain pharmacokinetic studies were conducted at the AAALAC accredited facility of Sai Life Sciences Limited (Pune, India). Briefly, male C57BL/6 mice were treated with a single intraperitoneal administration of BRD3290 at 3, 10, 30, or 56 mg/kg. Eighteen mice were used per dose group, with 3 mice per time point. The vehicle used for formulation was 5% v/v NMP, 5% Solutol HS-15 and 90% Captisol (20% v/v in reverse osmosis water). Blood samples (approximately 60 µL) were collected under light isoflurane anesthesia (Surgivet®) from retro orbital plexus from a set of 3 mice at 0.08, 0.5, 1, 2, 4, 8 and 24 hr post-injection. Immediately after blood collection, plasma was harvested by centrifugation at 4000 rpm for 10 min at 40 °C and samples were stored at -70 °C until bioanalysis. Following blood collection, animals were immediately sacrificed, and cerebrospinal fluid (CSF) was collected. The abdominal vena-cava was cut open and the whole body was perfused from the heart using 10 mL of normal saline. Brain samples were collected from a set of three mice at the same time point. After isolation, brain samples were rinsed three times in ice-cold normal saline and dried on blotting paper. Brain samples were homogenized using ice-cold phosphate buffer saline (pH 7.4). Total homogenate volume was 3 times the tissue weight. All homogenates were stored below -70°C until bioanalysis. All samples were processed for analysis by protein precipitation method and analyzed with fit-for-purpose LC-MS/MS method (Lower Limit of Quantification (LLOQ)= 1.02 ng/mL for plasma, 2.04 ng/mL for brain, and 5.12 ng/mL for CSF). The pharmacokinetic parameters were estimated using non-compartmental analysis tool of Phoenix® WinNonlin software (Ver 8.0).

### Cell line development and culture

Human AMPAR co-expression constructs consisted of codon-optimized paired tagged AMPAR subunits (Myc-*GRIA1i/o*, V5-*GRIA2i/o*, FLAG-*GRIA3i/o*) separated by a P2A auto-cleaving linker on a PCDNA5_FRT_TO backbone. Constructs were constructed at Genescript. Flip and flop isoforms were chosen for each subunit based on their prevalence in the human cortex and on their computationally predicted effect on the allosteric binding pocket. Q/R splice site editing was not recapitulated to allow the channels to pass calcium and enable a calcium-based fluorometric assay. Homomeric receptor constructs used architecture identical to that of heteromeric constructs, except they lacked the P2A linker and second subunit. A clonally selected Flp-In TREx-bearing HEK293T line with a stably integrated GCaMP6s-CAAX module, conferring Geneticin/G-418 resistance, was used as the parental cell line for all AMPAR cell lines. To create individual constructs, the parental cell line was plated at 10K cells/well in 24-well plates and transfected 48h after plating via Lipofectamine 2000 with 1 µg of the appropriate plasmid, grown for 3-5 weeks in 200 µg/mL hygromycin, 0.3 µg/mL Blasticidin and 10 µg/mL Geneticin/G-418, then replated at ∼1 cell/well in 24-well plates and expanded for an additional 2-3 weeks in selection antibiotics. Single colonies were visually identified and expanded for 4-8 weeks, validated via Western blot, coimmunoprecipitation, and sequencing. Final clones were selected based on growth rate and response in calcium flux FLIPR assay, and frozen in bulk using 10% DMSO in DMEM/F-12 with 10% FBS and 30 µM NBQX.

Cells were grown under 5% CO_2_ at 37C in DMEM/F-12 (Gibco# 10565-042) with 10% fetal bovine serum (FBS) (Gibco# 26140079) and 30 µM NBQX (Cayman# 14914-25), which prevented intrinsic cell death induced by the construct. Cells were kept within 20 passages of initial assessment in all assays and always kept under blasticidin and G-418 antibiotic selection except when plated in induction media for assays (<72h prior to assay). Cells were passaged twice a week once they reached 90-95% confluence. To passage, cells were washed once with room temperature (RT) PBS, then detached using Versene (Thermo Fisher Scientific). After 10min incubation at RT, Versene was neutralized with an equal amount of culture media, cells were centrifuged at 300g for 5 minutes, media was aspirated, and the cell pellet was resuspended in 10 mL of fresh culture media for counting, then plated at 0.1 million cells/mL in T75 or T175 flasks. Prior to any experiment, AMPAR construct expression was induced for 48h with 1 µg/mL doxycycline, and thawed cells were passaged at least three times before any cellular assay.

### Immunoblotting and immunoprecipitation

For immunoblotting, cells were plated at 1 million cells per well in a 6-well plate 72 hours prior to the planned experiment and AMPAR expression was induced with 1 µg/mL doxycycline. To extract cell lysate, plates were placed on ice, the media aspirated, and the cells were covered with 250 µL ice-cold PBS. Cells were scraped off the plate, washed with an additional 250 µL ice-cold PBS, and the plates were scraped a second time. The cell suspension was centrifuged for 5 minutes at 1,000 g at 4C, and PBS was subsequently removed. Cells were then lysed in 100 µL 1% Triton buffer (1% Triton, 1 mM EDTA, 50 mM Tris pH 7.5, 1:500 aprotinin, 1:200 leupeptin, 1:100 PIC2, 1:100 PIC3, 1:100 PMSF; milliQ water) for 15 minutes on ice. Lysate was centrifuged at 14,000 g for 10 minutes to pellet the insoluble fraction, and lysate was used for immunoblots. Protein concentration was normalized through BCA assay and 100 µL of protein was incubated for 16h at 4C on a rotator (VWR# 10136-084) with 1% of the appropriate pulldown antibody. At the end of this incubation, Protein A Dynabeads (ThermoFisher #10001D) magnetic beads were added to each tube, and the mixture was incubated on the nutator at 4C for an additional hour. Beads were magnetically collected, the unbound fraction removed for later analysis, and the beads washed three times with 150uL lysis buffer. Protein was liberated from the beads by addition of 50 µL 1x Laemmli buffer with 50 mM DTT. Maximal volumes of input, bound, and unbound fractions were mixed with 4x Laemmli buffer with 200 mM DTT and run on an 8-11% Bis-Tris Bolt gel (Thermo Fisher, #NW04127BOX) on ice for 15 minutes at 80V then 1h at 120V. Gels were transferred onto nitrocellulose membranes using iBlot transfer stacks (Thermo Fisher, #IB301001). All blocking and antibody incubation steps took place in 0.2 µm-filtered TBST (0.1% Tween-20) with 1% dry skim milk, and blots were washed three times following the removal of each antibody. Membranes were blocked for two hours on a shaker at room temperature, incubated with primary antibody at a dilution of 1:1000 for two hours at room temperature, incubated with secondary antibody at a dilution of 1:5000 for one hour at room temperature, and imaged on a ChemiDoc for 5-180s using SuperSignal Pico PLUS Chemiluminescent Substrate (Thermo Fisher# 34580). Immunoprecipitation blots were then stripped using 10 mL Restore PLUS Stripping Buffer (Thermo Fisher# 46430), re-blocked, and re-probed according to the above conditions.

### Immunocytochemistry

Cells were fixed at room temperature for 15 min with 4% paraformaldehyde (Electron Microscopy Sciences, Inc., Hatfield, PA) and 4% wt/vol sucrose (Sigma-Aldrich) in PBS (Sigma-Aldrich). After three washes with PBS, cells were permeabilized for 10 min at room temperature with 0.25% Triton-X100 in PBS and then washed twice with PBS. Following incubation with blocking buffer 5% BSA (Sigma-Aldrich) in PBS for 1 hr at room temperature, cells were then incubated with primary antibodies in blocking buffer overnight at 4 °C. After washing with PBS, the sample was incubated for 1 hr at room temperature with secondary antibodies and washed again three times with PBS. Lastly, neurons were incubated with DAPI for 5min and washed twice with PBS before imaging. Primary antibodies: Rabbit V5 (Cell signaling 13202s), mouse HA (Cell signaling 2367s), mouse Myc (Cell signaling 2276s)

### Electrophysiology

Automated patch clamp electrophysiology was performed using the Nanion Syncropatch 384-PE. Cells were plated at 5 million cells per T175 flask in 30 mL media containing 30 µM NBQX and cultured with 1 µg/mL doxycycline (Sigma D9891-10G) for 48h prior to the experiment. Cells were washed with room-temperature PBS prior to harvest, then harvested using Accutase (Millipore Sigma A6964). After 5 min of incubation, cells were triturated until the suspension consisted entirely of single cells as confirmed visually using a microscope. Accutase was then neutralized with an equivalent volume of media, followed by centrifugation of the suspension at 300g for 5 minutes, aspiration of the media, and resuspension of the cell pellet in 5 mL culture media. Centrifugation, aspiration and resuspension of the cell pellet in 5mL media was then repeated to ensure all Accutase was removed from the media. Cells were counted and diluted to a concentration of 1-2 million cells per mL in 12 mL of a 1:1 ratio of culture media without NBQX and external solution (140 mM NaCl, 5 mM glucose, 4 mM KCl, 2 mM CaCl_2_, 1 mM MgCl_2_, 10 mM HEPES; pH 7.4, osmolarity 295-305 mOsm). Cells were then placed in a Teflon reservoir, which was placed onto the Syncropatch “cell hotel” and shaken at 200 rpm at 10 °C with a lid to prevent evaporation. All experiments concluded within 2.5 h of initial cell dissociation to ensure cell health. Internal solution consisted of the following: 10 mM EGTA, 10 mM CsCl, 110 mM CsF, 10 mM HEPES, 10 mM NaCl; pH 7.2, osmolarity 280-290 mOsm. Cells were automatically loaded into a Nanion 384-well plate and the cells were captured by applying -140 mbar pressure for 30s or until >90% of wells displayed 0.1 MOhm seal quality. Seals were enhanced by applying a high-calcium seal enhancer (140 mM NaCl, 5 mM glucose, 4 mM KCl, 10 mM CaCl_2_, 1 mM MgCl_2_, 10 mM HEPES; pH 7.4, osmolarity 295-305 mOsm) in the external chamber to introduce cesium fluoride crystallization around captured cells, yielding gigaohm seals necessary for high-quality patch clamp recordings. Cells were then washed 3 times to reduce the extracellular Ca^2+^ level back to 5 mM CaCl_2_, and all recordings were performed in external recording solution (140 mM NaCl, 5 mM glucose, 4 mM KCl, 2 mM CaCl_2_, 1 mM MgCl_2_, 10 mM HEPES; pH 7.4, osmolarity 295-305 mOsm). Two transient applications of -200 mbar pressure enabled whole-cell recordings, with cells returned to -50 mbar holding pressure and voltage clamped at -80 mV for the remainder of the recording.

Compounds were resuspended in high-calcium external solution (140 mM NaCl, 5 mM glucose, 4 mM KCl, 5 mM CaCl_2_, 1 mM MgCl_2_, 10 mM HEPES; pH 7.4, osmolarity 295-305 mOsm) in a seven-point half-log dilution series ranging from 100 µM to 100 nM. The DMSO concentration was adjusted to 0.5% in all compound additions and compound-free control recordings. AMPAR currents were recorded during three separate 500 ms quick liquid additions (aka ligand puffs): once in response to high-calcium external recording solution alone, once in response to high-calcium external recording solution containing 3 mM glutamate, and once in response to high-calcium external recording solution containing 3 mM glutamate following three minutes of PAM incubation. Puffs were delivered as stacked solutions, loading 20 µL of high-calcium external solution followed by 5 µL of 3 mM glutamate in high-calcium external solution, delivered to the wells at 10 µL/s for 2.5s. The wells were washed with standard (140 mM NaCl, 5 mM glucose, 4 mM KCl, 2 mM CaCl_2_, 1 mM MgCl_2_, 10 mM HEPES; pH 7.4, osmolarity 295-305 mOsm) external recording solution between all puffs to prevent crossover. Liquid handler tips were washed between additions with 2 mM EDTA in sterile RO H_2_O instead of ethanol, as ethanol was found to disrupt the temporal precision of puff delivery, likely by interfering with compound stacking. Current density was calculated by taking the peak current during compound + glutamate addition and dividing by the capacitance of the cell. Cell response to glutamate alone in the absence of compound was used to measure overall cell quality over time and between runs.

Any low-quality cells that met one or more of the following exclusion criteria were excluded from analysis: wells that displayed seal resistance less than 1 MOhm or greater than 100 MOhm, cell capacitance below 5 pF or above 200 pF, or series resistance below 1 MOhm or above 40 MOhm. Each data point from each concentration of each compound on each day represents an average of at least three electrodes’ recordings. All cell and compound testing locations were alternated on the plate between runs to minimize any plate or electrode-dependent effects. Concentration-response curves were constructed in Prism and compared to within-plate DMSO only control wells.

### High throughput fluorescence assay

Cells were plated 2-3 days prior to the experiment at 10,000 cells/well in 50 µL/well of culture media (DMEM/F-12 with GlutaMax, 10% FBS, 30 µM NBQX) with 1µg/mL doxycycline onto black poly-d-lysine-coated 384-well clear-bottom plates (Corning) and were roughly 80% confluent at the time of recording. On the day of the assay, compound in DMSO was added into high-calcium FLIPR buffer (10mM HEPES, 140mM NaCl, 5mM KCl, 1mM MgCl_2_, 15 mM CaCl_2_; pH 7.3) in 384w compound plates at a dilution of 1:200 using the CyBio Well Vario Pipettor (Analytik Jena). Immediately prior to recording, cells were washed twice with FLIPR assay buffer (10mM HEPES, 140mM NaCl, 5mM KCl, 1mM MgCl_2_, 0.5 mM CaCl_2_; pH 7.3) to remove the growth media and maintained in 15 µL of assay buffer for less than 10 minutes at RT prior to recording. Data was recorded using the Molecular Devices FLIPR Penta recording at 1Hz with changes in fluorescence measured over time with an excitation at 470-495 nm and emission at 515-575 nm and the recording chamber maintained at 25 °C. Baseline data were collected first for 30 s, followed by an addition of 15 µL of compound in high-calcium (15 mM CaCl_2_) assay buffer and recorded for 180s to observe agonist activity. This was followed by a subsequent addition of 15 µL assay buffer containing 120 µM glutamate, then recorded for another 180s to measure PAM activity. The final addition of 15 µL of assay buffer containing 3 µM BIIB-104 was used to establish a antagonist activity, and was recorded for an additional 180s. Compounds were assayed in 10-point dose response, with technical duplicates across three independent biological replicates. Internal PAM, agonist, and inhibitor controls were included on every plate in concentration-response curve. Compounds were assayed for EC_50_ and E_max_, with known AMPAR PAM BIIB-104 (3 µM) response establishing 100% activity, an equivalent concentration of DMSO as the neutral control, and the selective AMPAR inhibitor NBQX establishing 0% activity on each plate.

### FLIPR and Automated Patch Clamp Electrophysiology Data Normalization Methods

From the FLIPR data, we utilized an activity score to generate a single value that utilizes both the E_max_ and the EC_50_ of each compound for each cell line based on previous work ^19^.

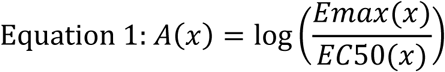

To create an assay-independent value that could be used to compare compound activity across cell lines, all FLIPR activity data was normalized against HBT1, which was selected as our reference compound as it behaved most consistently in our hands when compared to other reported compounds and displayed no intrinsic agonist activity across the various cell lines tested. Using the FLIPR data, we then calculated D_(cell line,cpd)_ as the uncorrected delta activity: the activity score of a compound in a given cell line, minus the activity score of HBT1 in the same cell line, which was collected within the same assay plate.

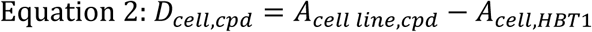

We then set out to characterize the selectivity of HBT1 across the eight cell lines used in this study to generate a correction factor that can be used to correct for the inherent selectivity of HBT1 across the eight cell lines. A single receptor type (GluA1i/2o) was chosen as the reference activity level. For the other receptor types, the difference in activity score compared to GluA1i/2o was taken as a "correction factor". This created eight distinct correction factors, one for each cell line.

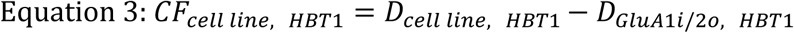

The corrected Δlog is calculated by applying experimentally derived correction factors (CF) based on 26 published nonselective AMPAR PAMs. Each cell line has its own correction factor. The Δlog _cell line, cpd_ for a given compound tells us how active this compound is compared to HBT1 in a given cell line, after correcting for the intrinsic selectivity of HBT1.

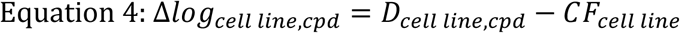

ΔΔ*log_cell line_*_,*cpd*_ then describes the selectivity of a compound for a given cell line compared to a reference cell line (“ref”) – always GluA1i/2o in our analysis.

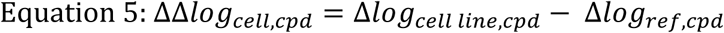

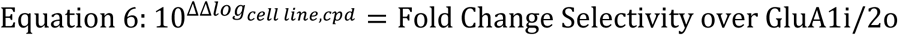

The same process is then applied to automated patch clamp electrophysiology data, with one key difference: HBT1 data is still collected on the same day for a given cell line but using a separate assay plate from the test compound. This is because automated patch clamp electrophysiology has lower throughput than the FLIPR assay.

### cGMP biomarker assay

Adult male CD-1 mice from Charles River Laboratories aged between P80 and P100 weighing 25-35g were dosed intraperitoneally (i.p.) with compound or vehicle 30 minutes prior to tissue collection. Animals were randomly assigned to each group and each group consisted of 8-18 mice. For PF4778574, animals were dosed at 1mg/kg at a volume of 10mL/kg i.p. Vehicle consisted of 5:5:90 (v/v/v) DMSO:Cremophor EL:deionized H_2_O. Effects of BRD3290 on cGMP was determined using a similar dosing paradigm as PF-4778574 but dosed at 56 mg/kg at a volume of 10mL/kg i.p. in 5:5:90 (v/v/v) NMP:Solutol-HS15:Captisol (20% w/v in RO water). 30min after dosing compound or vehicle, mice were euthanized via rapid cervical dislocation (no anesthesia) and the cerebellum rapidly transferred to pre-weighed Eppendorf tubes and snap frozen on liquid nitrogen. Tubes were then subsequently weighed and the total cerebellum tissue calculated. cGMP was extracted from cerebellar tissue samples and measured with a cyclic GMP ELISA kit (Cayman Chemical # 581021) following kit directions. Briefly, cerebellar brain tissue was lysed in 10 mL of 5% trichloroacetic acid (TCA) per mg of brain tissue and homogenized via a tip sonicator (20s; power setting 3; 50% activation). TCA was removed from samples via triplicate organic/aqueous phase extractions with 5 volumes of water-saturated diethyl ether (Millipore Sigma# 309966-1L). The remaining diethyl ether was removed by evaporation at 70 °C for 10 minutes, which we found to provide more reliable results than the 2-minute evaporation described in the kit. No acetylation of the sample was required when samples were diluted 1:50 in ether-extracted 5% TCA, and standards were prepared in a two-fold dilution curve in ether-extracted 5% TCA according to kit instructions. 50 µL of included ELISA buffer was added to NSB wells, and 50 µL of ether-extracted 5% TCA was added to background non-specific binding (NSB) wells and maximal binding wells. 50 µL of diluted sample was added to sample wells, with each sample measured in technical triplicate, and 50 µL of the appropriate standard was added to each standard well, which were assayed in technical duplicate as per kit instructions. 50 µL of cGMP-acetylcholinesterase conjugate tracer was added to each well except the blank and tracer-alone wells. 50 µL cGMP ELISA antibody was added to all wells except the blank, tracer-alone, and NSB wells. This plate was covered with included plastic film and incubated overnight for 18 hr at 4 °C. The following day, immediately prior to assay, Ellman’s reagent was reconstituted. Wells were emptied, washed five times with included wash buffer, and 200 µL of Ellman’s reagent was placed in each well. 5 µL of cGMP-acetylcholinesterase tracer was added to the tracer-alone well on each plate, the plate was covered with a new sheet of included plastic film, and plates were incubated for 2 hr in the dark at room temperature on an orbital shaker at 200 rpm. At the end of the incubation, the plate was cleaned and read on a Perkin Elmer Enspire MultiMode Plate Reader (MLD2300_0000) at 410 nm. Data were processed using the calculator worksheet included with the kit, and cGMP response was then normalized to that of vehicle-treated mice for each compound. Kits were used within four weeks of reconstitution as per the manufacturer’s recommendations.

### Convulsion Liability

Male C57BL/6J mice, aged 3-5 months were used and were not handled prior to testing. Briefly, mice were habituated to the test room in their home cages for 60 minutes during the light cycle. For the PF4778574 study, mice were weighed and then dosed with vehicle (5:5:90 (v/v/v) DMSO:Cremophor EL: RO H2O), 1 or 1.78mg/kg PF4778574 subcutaneous (s.c.) at 10mL/kg. For the BRD3290 study, mice were dosed with vehicle 5:5:90 (v/v/v) NMP:Solutol-HS15:Captisol (20% w/v in RO H_2_O) or 10, 30 or 56 mg/kg BRD3290 i.p. Mice were then placed into clean cages for observation. The mice were monitored continuously for 3 hours post-injection and video recorded. Mice were monitored for seizure-like symptoms using a modified Racine scale^22^. Mice that showed seizure-like symptoms were immediately euthanized via cervical dislocation. Mice that did not display convulsions were returned to their home cages following the 3-hour test period.

### Accelerating Rotarod

Mice were not handled prior to testing and were habituated to the testing room in their home cages for 30 minutes prior to testing in white light. The first day of testing consisted of multiple training phases. First, the mice were placed onto the stationary rotarod apparatus (Harvard Apparatus LE8200) for 1 minute. Then the mice had a minimum of two trials with the rotarod rotating at a constant 4 RPM (no acceleration). For the first trial, mice were put back onto the rod if they fell off before 1 minute or removed if they remained on the rod for 1 minute, whichever came first. For the second trial, mice were put back onto the rod if they fell off in under 1 minute. Mice that remained on the rod for 1 minute completed training. Mice that did not stay on the rod for 1 minute were given up to 2 more trials. Mice that could not stay on the 4 RPM rotating rod for 1 minute by the 4th trial were excluded from further testing. At the end of training, the mice were returned to their home room. The apparatus was cleaned with 70% ethanol in between trials.

24 hr after Rotarod training, mice were returned to the test room and allowed to habituate to the room for 30 minutes. Mice were placed on the rotating rod, and acceleration was started once all mice were on the rod. Acceleration gradually increased from 4 to 40 RPM over 60 seconds. The time it took for the mice to fall off the rotating rod was recorded, and mice were then returned to their home cage. Mice were tested 3 times with a minimal 5-minute intertrial interval. The apparatus was cleaned with 70% ethanol in between trials. For the PF4778574 study, mice were dosed 30 minutes before testing with PF4778574 (0.56mg/kg) or vehicle (5:5:90 (v/v/v) DMSO:Cremophor EL: RO H_2_O) s.c. at 10mL/kg. For the BRD3290 study, mice were dosed with BRD3290 (3, 10, 30, or 56 mg/kg) or vehicle (5:5:90 (v/v/v) NMP:Solutol-HS15:Captisol (20% w/v in RO H_2_O) i.p. All experiments were performed during the light cycle.

### Spontaneous Alternation

Mice were habituated to the test room, in white light, for 30 minutes prior to dosing. Mice were first dosed with PF-4778574 (0.032, 0.056, 0.1, or 0.32mg/kg) or vehicle (5:5:90 (v/v/v) DMSO:Cremophor EL: RO H_2_O) s.c. 30 minutes prior to initiation of the assay. 10 minutes after administration of PF-4778574 or vehicle, mice were dosed with MK801 (0.18mg/kg) or 0.9% saline i.p. 20 minutes after MK-801 or saline injection, mice were placed into one arm of Y-maze (each arm is 39 cm long) and allowed to freely explore the entire apparatus for 8 minutes. The starting arm was alternated for each mouse and was randomly assigned prior to testing. The trials were recorded using Ethovison 17. The apparatus was cleaned with 70% ethanol after each trial. All experiments were performed during the light cycle. The number and order of entries into each of the three arms were analyzed. A correct spontaneous alternation occurred when the mouse entered a different arm in each of three consecutive arm entries (e.g., ABC or CAB is correct, ABA or CAC is incorrect). Percent alternation was calculated as:

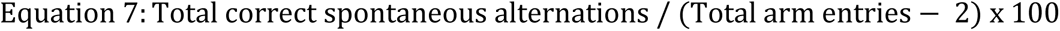

### Novel Object Recognition

Mice were handled for three days prior to testing. Mice were first habituated to the testing room in their home cages for 30 min prior to dosing. To acclimate the mice to the testing chamber, they were added to a clean and empty Ethovision open field box (40 × 40 cm) and allowed to freely explore the apparatus for 15 minutes under white light. The mouse was then removed, placed back into its home cage and returned to the home room and the box was cleaned with Clidox® (Pharmacal Research Laboratories). 24 hr after habituation to the testing chamber, mice were brought back to the testing room and habituated for 30 minutes in white light. Mice were then dosed with either PF-4778574, BRD3290, or vehicle 30 minutes prior to the start of the familiarization phase. Two sets of objects were used, 2 tall stacks of Legos® or 2 T75 flasks filled with used desiccant. These objects were confirmed to have comparable salience using a separate cohort of WT mice. During the familiarization phase, the open field box contained two identical objects, with one placed in the upper left-and one in the upper right-hand corners of the arena. The mouse was then placed into the arena containing two identical objects and allowed to explore the objects for a total of 20 seconds or until the mouse was in the box for 10 minutes, whichever came first. The mouse was then removed, and the box and objects were cleaned with Clidox® between trials. 24 hr after the familiarization phase, the mice were returned to the test room and habituated for 30 minutes prior to dosing. Mice then received the same compound and dose as the day prior. Before placing mice in the arena, one of the previously explored objects was replaced with a novel object (Legos or flask). 30min after dosing, the mouse was then placed into the box and allowed to freely explore for 10 minutes. The time spent exploring each object was quantified automatically using Ethovision 17. For the PF4778574 studies, mice were dosed with PF4778574 (0.032, 0.056, 0.1, or 0.32mg/kg) or vehicle s.c. For the BRD3290 study, mice were dosed i.p. with either BRD3290 (3, 10, 30, or 56 mg/kg) or vehicle. All animals were dosed 10mL/kg body weight, and all experiments were performed during the light cycle of the animals under white light.

## Supporting Information

### Author Contributions

The manuscript was written through contributions of all authors. All authors have given approval to the final version of the manuscript.

### Funding Sources

This work was supported by the Stanley Center for Psychiatric Research, at the Broad Institute of MIT and Harvard

## ACKNOWLEDGMENT

We would like to thank the Stanley Family Foundation’s tremendous support of the Stanley Center since its inception in 2014. We would like to also thank Dr. Yuemin Bian for his help with data analysis and acknowledge Godhuli Nandi, Suman Kumar Joardar, and Abhijit Kundu for their valuable contributions to the project’s synthetic chemistry.

## ABBREVIATIONS

AMPA: α-amino-3-hydroxy-5-methyl-4-isoxazolepropionic acid
AMPAR: AMPA receptor
CSF: cerebrospinal fluid
DMPK: drug metabolism and pharmacokinetics
FDA: Food and Drug Administration
i.p.: intraperitoneally
LLOQ: Lower Limit of Quantification
NOR: Novel Object Recognition
NSB: Nonspecific Binding
PAM: Positive Allosteric Modulator
S.C.: subcutaneous
TCA: Trichloroacetic Acid

## Supporting Information

**Supplemental Figures and Methods:** Cell line validation, extensive pharmacokinetic characterization of BRD3290, effects of AMPAR PAM in spontaneous alternation and compound synthesis methods.

**Supplemental Tables S1-S4:** Supplemental Tables that contain the extended FLIPR data for key SAR molecules and the in vitro ADME properties of BRD3290.

## Notes

### Competing Interest Statement

The authors have declared no competing interest.

### Summary of Updates

Corrected small error in one of the chemical structures

