## Supplemental Figures and Materials for "Discovery of GluA3 preferring AMPA receptor positive allosteric modulator BRD3290"

**Supplemental Materials and Methods**

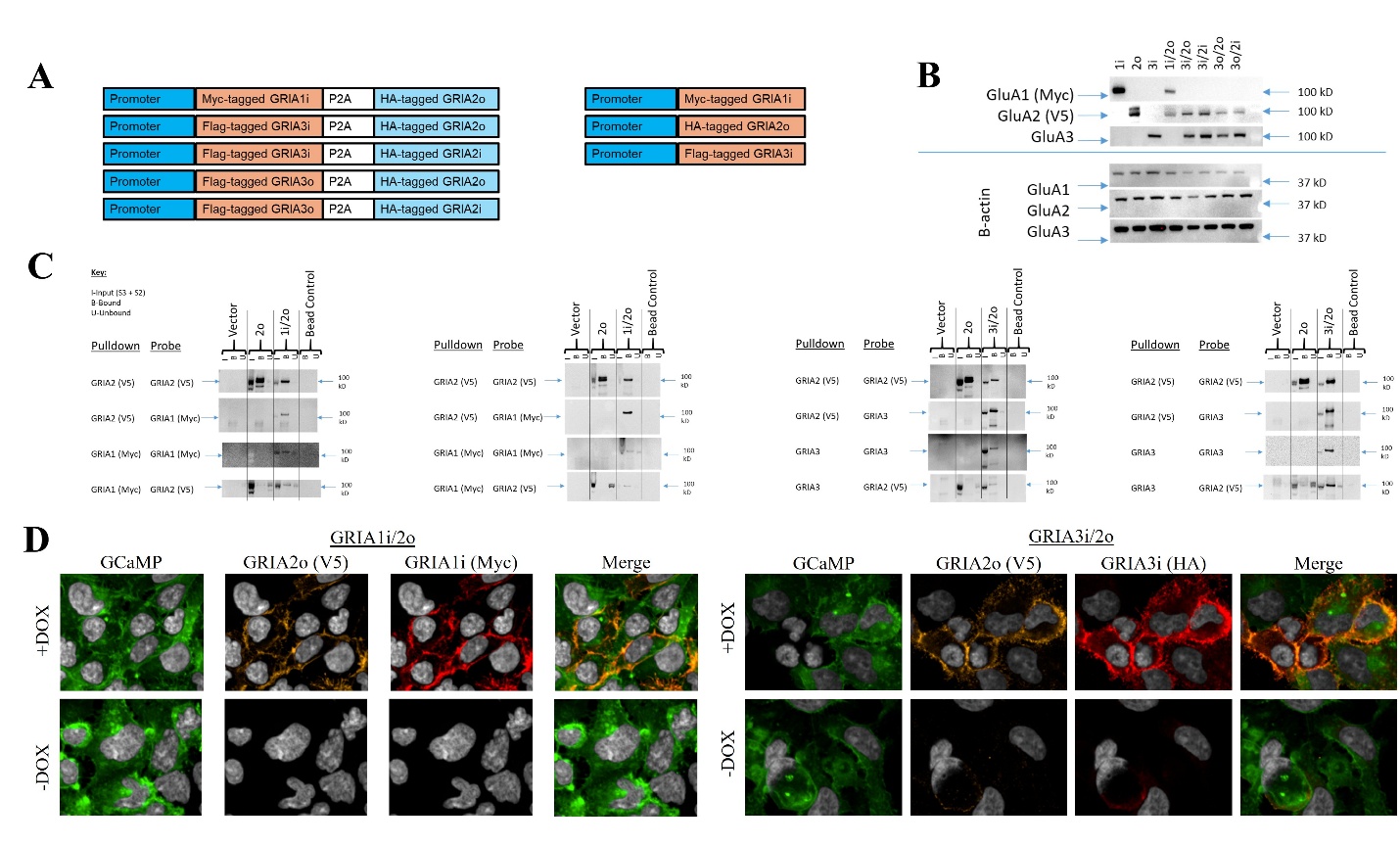
**Supplemental Figure 1:** AMPAR cell line construction and validation (A) Diagram of AMPAR homodimer/heterodimer and GCaMP6s-CAAX reporter constructs. (B) Western blot of all eight clonal cell lines. (C) Co-immunoprecipitation and (D) immunocytochemistry imaging of AMPAR heterodimers in primary target (GluA3i/2o) and anti-target (GluA1i/2o) cell lines in induced (+DOX) and basal (-DOX) conditions.

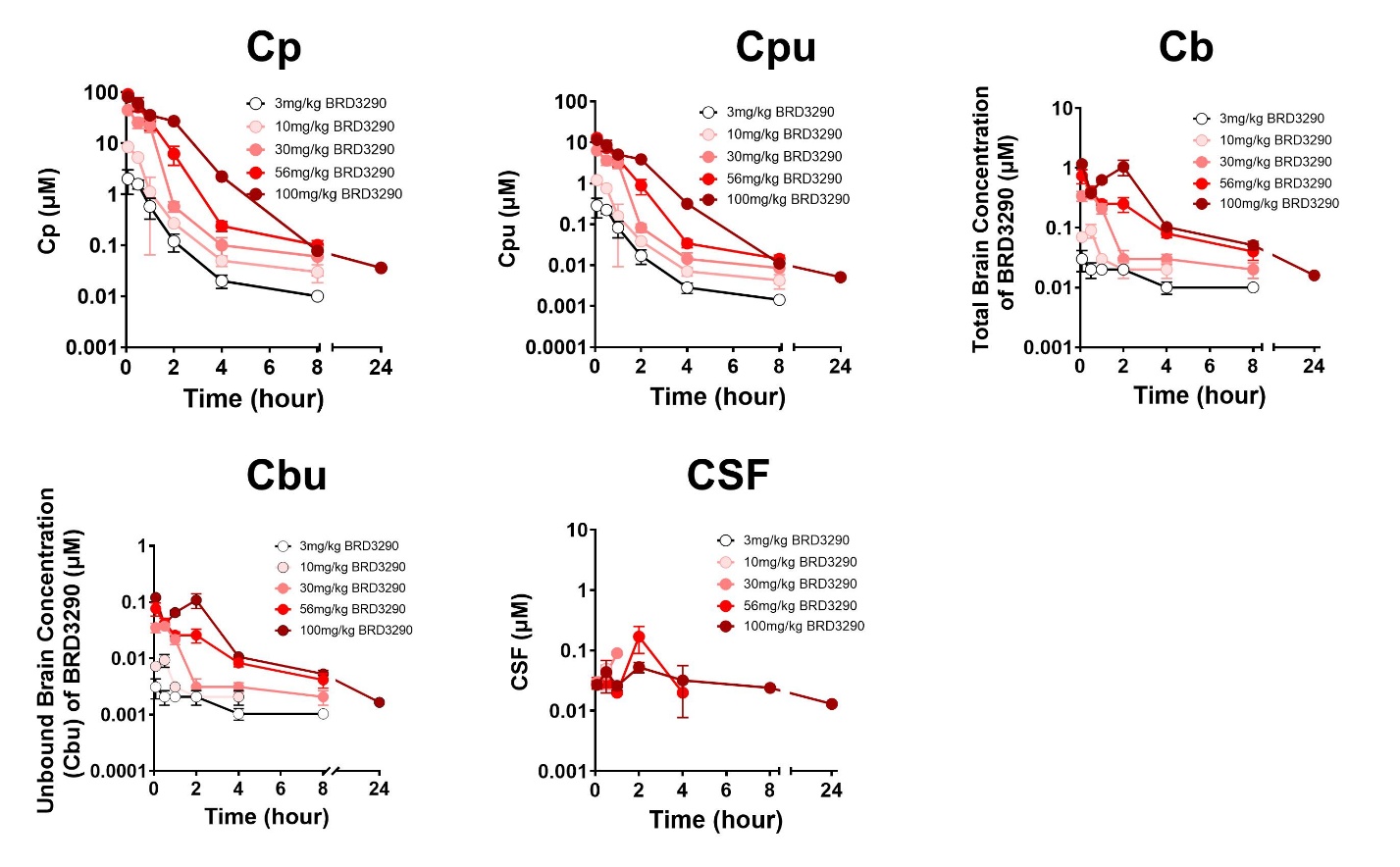

**Supplemental Figure 2**: Comprehensive pharmacokinetic dataset of BRD3290 in C57BL/6 mice. (A) Plasma concentration (C_p_), unbound plasma concentration (C_p,u_), brain concentration (C_b_), unbound brain concentration (C_b,u_), and CSF concentration (CSF) of BRD3290 in mice dosed i.p. with BRD3290. Data points display mean ± SEM of n = 3 mice per timepoint per dose. Lower limit of quantification = 1.02 ng/mL for plasma, 2.04 ng/mL for brain and 5.12 ng/mL for CSF; data points below LLOQ not displayed for clarity, though data collected for all doses at all timepoints.

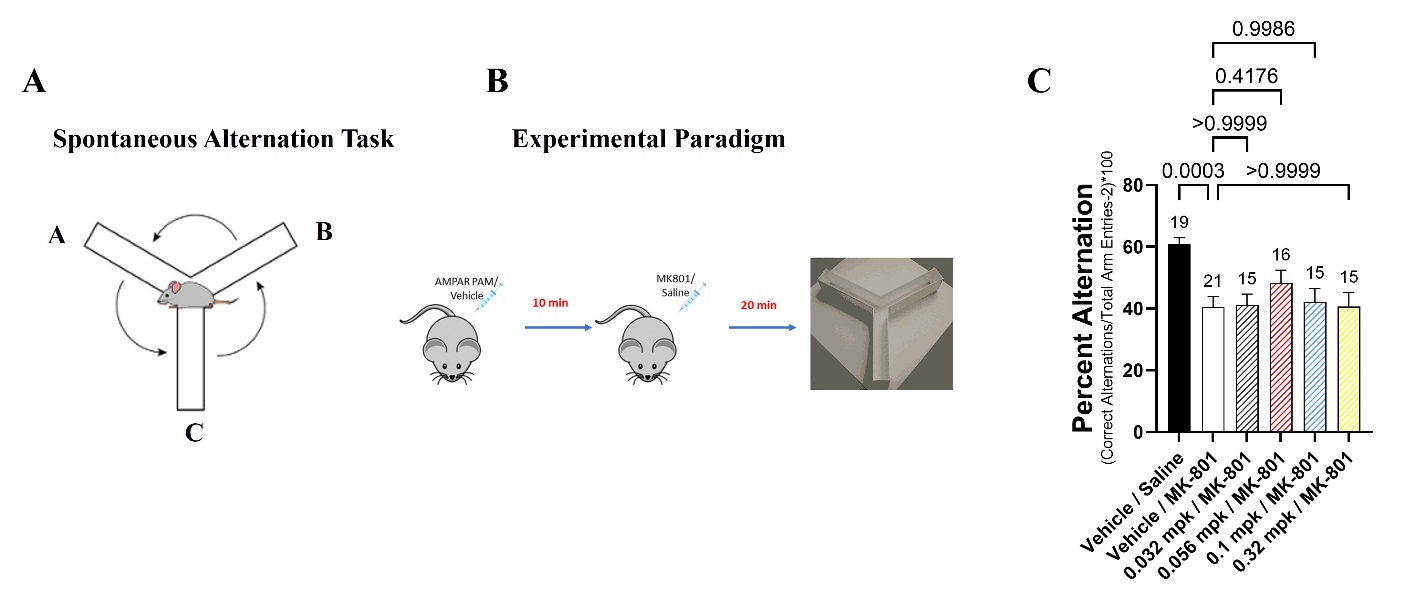
**Supplemental Figure 3:** AMPAR PAM PF-4778574 does not reverse MK-801 induced deficits in spontaneous alternation. (A) Diagram of spontaneous alternation task and (B) MK-801 induced deficit paradigm in rodent spontaneous alternation task. (C) Percent correct alternation; Data points display mean ± SEM. One-way ANOVA with Dunnett’s post hoc test. Number of mice per group displayed above each bar.

**Compound synthesis section**

**General methods:**

Reagents and solvents were obtained from commercial vendors and used as received.

^1^H NMR spectra were recorded on a Bruker 400.32 MHz Ultra Shield, on Avance Neo spectrometer. Chemical shifts (δ, ppm) are reported relative to the solvent peak (DMSO-d6: 2.50 [1H]). Proton resonances are annotated as: chemical shift (δ), multiplicity (s, singlet; d, doublet; t, triplet; q, quartet; m, multiplet; br, broad), coupling constant (J, Hz), and number of protons.

^13^C NMR spectra were recorded on a Bruker Avance Neo 600 MHz equipped with a 5mm helium-cooled QCI-F cryoprobe.

Liquid chromatography–mass spectrometry (LC–MS) analysis was carried out on a Waters ACQUITY H-Class Ultra Performance Liquid Chromatography (UPLC) system coupled with a Waters SQD 2 Mass Spectrometer. Mass spectrometric detection was performed using an Electrospray Ionization (ESI) source.

High resolution mass spectrometry (HRMS) was recorded on an Agilent 6545 QTOF mass spectrometer coupled to an Agilent Infinity 1260 LC system running a Jet Stream electrospray ionization (ESI) source.

Specific optical rotation was measured on an Anton Paar MCP 200 polarimeter using a 100.00 mm path-length cell at a sample concentration of 0.2533 g/100 cm³.

All solvents were purified by standard drying procedures prior to use. Tetrahydrofuran was distilled from sodium and benzophenone under a nitrogen atmosphere. Dichloromethane and toluene were distilled from calcium hydride under nitrogen and stored over activated molecular sieves.

**Synthesis of BRD3290**

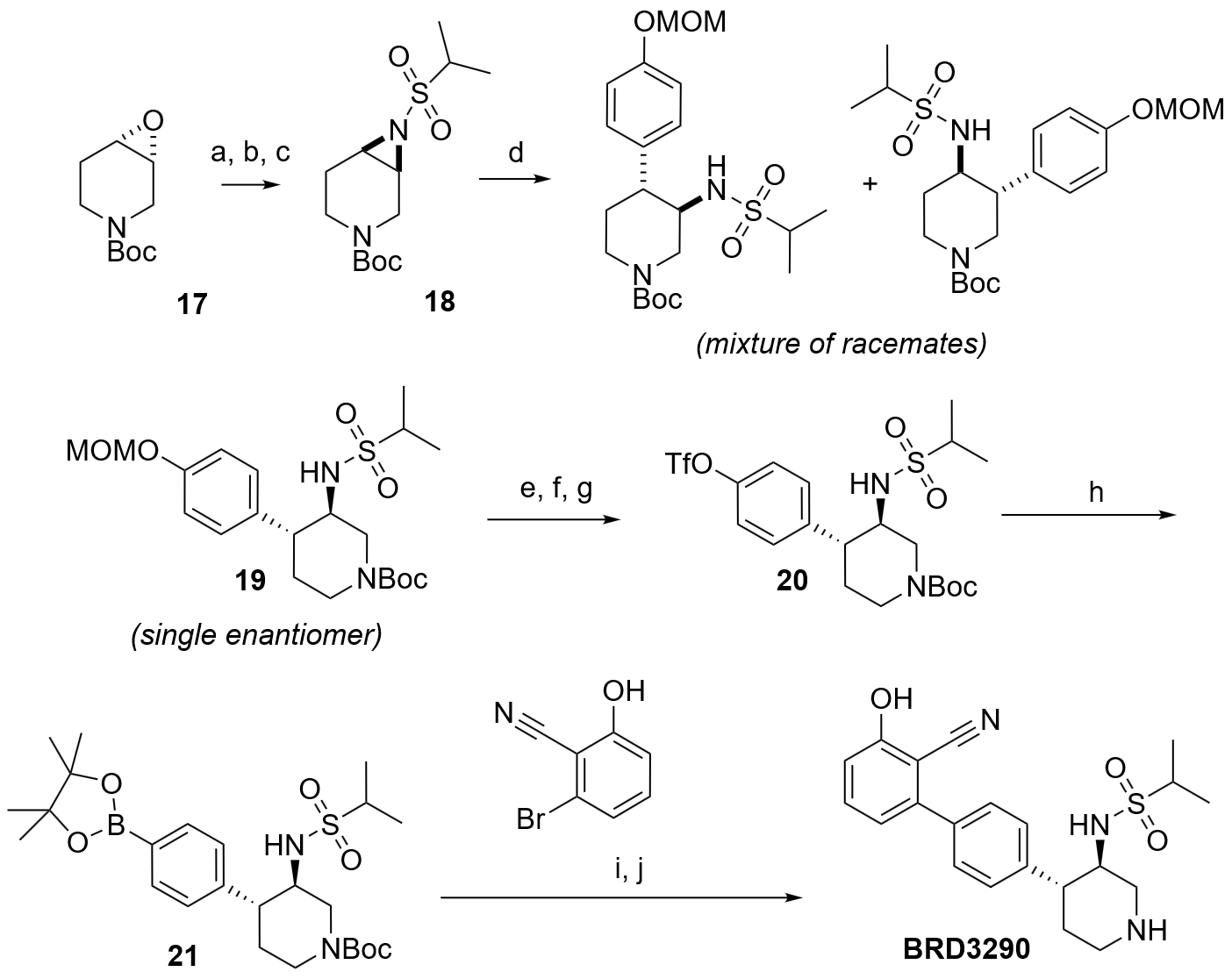

Reagents and conditions: (a) iPrSO_2_NH_2_, K_2_CO_3_, TEBAC, dioxane (b) MsCl, pyridine, DCM (c) K_2_CO_3_, MeCN; (d) ArMgBr, Cu(OTf)_2_, THF; (e) HCl, dioxane (f) Boc_2_O, Et_3_N, MeOH (g) Tf_2_O, pyridine, DCM; (h) bis-pin, KOAc, Pd(dppf)Cl_2_, dioxane; (i) ArBr, K_3_PO_4_, Pd(PPh_3_)_4_, dioxane, H_2_O (j) HCl, dioxane.

**Step a: Synthesis of tert-butyl (3R,4R)-3-hydroxy-4-((1-methylethyl) sulfonamido) piperidine-1-carboxylate &** **tert-butyl (3S,4S)-4-hydroxy-3-((1-methylethyl) sulfonamido) piperidine-1-carboxylate**

To a stirred solution of epoxide **17** (40 g, 201 mmol, 1.00 eq) in dioxane (100 mL), K_2_CO_3_ (27.75 g, 201 mmol, 1.00 eq), TEBAC (13.72 g, 60.2 mmol, 0.300 eq) and propane-2-sulfonamide (29.67 g, 241 mmol, 1.20 eq) were added and the reaction mixture was heated to 85 ºC for 16 h. After the completion of the reaction (checked by TLC; 50% ethyl acetate-hexane solvent system, new spot R_f_ 0.3; SM R_f_ 0.5), the reaction mixture was extracted with ethyl acetate, washed with water, brine, dried over Na_2_SO_4_ and concentrated to afford the crude compound.

The crude was purified by column chromatography using ethyl acetate-hexane solvent to afford the desired compound as a mixture of regioisomers (20 g, 62.0 mmol, 31% yield)

ESI m/z calculated: 222.1, found: 223.2 [MH+]

**Step b: Synthesis of** **tert-butyl (3R,4R)-4-((1-methylethyl) sulfonamido)-3-((methylsulfonyl) oxy) piperidine-1-carboxylate &** **tert-butyl (3S,4S)-4-((1-methylethyl) sulfonamido)-3-((methylsulfonyl) oxy) piperidine-1-carboxylate**

The mixture of isomers from the previous step (40 g, 124 mmol, 1.00 eq) was dissolved in DCM (300 mL) and cooled to 0 ºC in an ice-water bath. Pyridine (50 mL, 620 mmol, 5.00 eq) was added dropwise to the solution and the mixture was stirred for 20 minutes. Methanesulfonyl chloride (67 mL, 620 mmol, 5.00 eq) was added dropwise to the solution and the mixture was allowed to warm to room temperature and stirred for 16 h, after which consumption of starting material was observed by LC-MS.

The reaction mixture was washed with 1M HCl, and the aqueous layer was re-extracted with ethyl acetate. The organics were combined, washed with brine and dried over magnesium sulfate. The solvent was removed under vacuum to afford the crude mixture of mesylate regioisomers (32 g, 79.9 mmol, 64% yield) which was submitted to the next step with no further purification or characterization.

**Step c: Synthesis of tert-butyl (1R,6S)-7-(isopropylsulfonyl)-3,7-diazabicyclo[4.1.0]heptane-3-carboxylate (18)**

To a stirred solution of a mixture of crude mesylates from the previous step (20 g, 50 mmol) in acetonitrile (150 mL), K_2_CO_3_ (8.6 g, 62.4 mmol, 1.25 eq) was added and the reaction mixture was stirred at room temperature for 16 h.

After the completion of the reaction (checked by TLC; 50% ethyl acetate/hexane), the reaction mixture was extracted with ethyl acetate, washed with water, brine, dried over Na_2_SO_4_ and concentrated to afford the crude compound.

The crude was purified by silica column chromatography using ethyl acetate/hexane to afford the product (10 g, 32.9 mmol, 66% yield) as an oil.

ESI m/z calculated: 304.2, found: 305.2 [MH+]

**Step d: Synthesis of tert-butyl rac-(3R,4R)-3-(isopropylsulfonylamino)-4-[4-(methoxymethoxy)phenyl]piperidine-1-carboxylate (19)**

Tert-butyl rac-(1S,6R)-7-isopropylsulfonyl-3,7-diazabicyclo[4.1.0]heptane-3-carboxylate (10 g, 32.9 mmol, 1.00 eq) was dissolved in THF (200 mL) and the solution was sparged with argon for 15 minutes. Copper(II) trifluoromethanesulfonate (1.19 g, 3.29 mmol, 0.100 eq)  was added. The mixture was cooled to -78 ºC and stirred for 30 minutes. (4-(methoxymethoxy)phenyl)magnesium bromide (23.8 g, 98.6 mmol, 3.00 eq as a solution in THF) was added dropwise and stirring was continued overnight, allowing the mixture to warm to room temperature. The reaction mixture was cooled in an ice-water bath and quenched with aqueous NH_4_Cl solution. The mixture was diluted with ethyl acetate, washed with brine, and dried over magnesium sulfate and purified by silica chromatography (70% ethyl acetate in hexane) to afford a mixture of regioisomers (13 g) as an off white solid.

The mixture was further purified by preparative SFC (Column Name: CHIRALPAK IC, Solvent CO_2_ + 0.3% isopropylamine in MeOH) to afford the desired single enantiomer of compound **19** (5.7 g, 12.9 mmol, 39%).

^1^H NMR (400 MHz, DMSO-*d_6_*): δ 7.24 (d, J = 8.4Hz,2H), 7.05 (d, J = 8.8Hz,1H), 6.94 (d, J = 8.4Hz,2H), 5.14 (s, 2H), 4.29 (s, 1H), 3.95 (m, 1H), 3.33 (s, 3H), 3.16 (m,1H), 2.39 (m, 1H), 1.71-1.61 (m, 2H), 1.41 (s, 9H ), 1.33 (m,1H), 0.95 (d, J = 6.8 Hz, 3H), 0.50 (d, J = 6.6 Hz, 3H) ppm­­­.

ESI m/z calculated: 442.2, observed 443.2 [MH+]

Chiral purity = 99.94% ee (RT = 2.73 min) [Chiral Pak IC].

**Steps e and f: Synthesis of** **tert-butyl (3R,4R)-4-(4-hydroxyphenyl)-3-((1-methylethyl) sulfonamido) piperidine-1-carboxylate**

To a stirred solution of tert-butyl (3R,4R)-3-(isopropylsulfonylamino)-4-[4-(methoxymethoxy) phenyl] piperidine-1-carboxylate (750 mg, 1.69 mmol, 1.00 eq) in dioxane (3ml) was added HCl (4 M in dioxane, 4.24 mL, 16.9 mmol, 10.0 eq) and the reaction mixture was stirred for 4h at room temperature. After complete consumption of starting material, solvent was removed under reduced pressure, and the crude material was dissolved in dioxane (7 ml) and MeOH (7 mL). Triethylamine (1.43 mL,10.2 mmol, 6.00 eq) was added followed by boc anhydride (0.39 mL, 1.69 mmol, 1.00 eq) and the mixture was stirred for 16 h at room temperature. The solvent was evaporated, and the crude was purified by silica chromatography (50% ethyl acetate-hexane) to afford tert-butyl (3R,4R)-4-(4-hydroxyphenyl)-3-(isopropylsulfonylamino) piperidine-1-carboxylate (550 mg, 81% yield)

 ESI m/z calculated: 398.2, Observed 399.2 [MH+]

**Step g: Synthesis of tert-butyl (3R,4R)-3-((1-methylethyl) sulfonamido)-4-(4-(((trifluoromethyl)sulfonyl)oxy)phenyl)piperidine-1-carboxylate (20).**

Tert-butyl (3R,4R)-4-(4-hydroxyphenyl)-3-(isopropylsulfonylamino) piperidine-1-carboxylate (450 mg, 1.13 mmol, 1.00 eq) was dissolved in DCM (5 mL) and Pyridine (5 mL) and trifluoromethanesulfonic anhydride (0.28 mL, 1.69 mmol, 1.50 eq) was added dropwise at 0°C. The reaction was stirred for 1.5 h with gradual increase of temperature to room temperature. After completion of the reaction (monitored by TLC) the mixture was evaporated and purified by silica chromatography (60% ethyl acetate-hexane), affording tert-butyl (3R,4R)-3-(isopropylsulfonylamino)-4-[4-(trifluoromethylsulfonyloxy) phenyl] piperidine-1-carboxylate **4** (450 mg,75% yield).

^1^H NMR (400 MHz, DMSO-*d_6_*): δ 7.55 (d, J = 8.4Hz,2H), 7.43 (d, J = 8.8Hz,2H), 7.14 (d, J = 8.4Hz,1H), 4.34 (s, 1H), 4.00 (d, 1H), 3.27 (m, 1H), 2.68 (m, 3H), 2.49 (m,1H), 1.73 (m, 2H), 1.41 (s, 9H ), 1.33 (m,1H), 0.93 (d, J = 6.8 Hz, 3H), 0.48 (d, J = 6.6 Hz, 3H) ppm­­­.

 ESI m/z calculated: 530.1, observed 531.2 [MH+]

**Step h: Synthesis of tert-butyl (3R,4R)-3-(isopropylsulfonylamino)-4-[4-(4,4,5,5-tetramethyl-1,3,2-dioxaborolan-2-yl) phenyl] piperidine-1-carboxylate (21)**

Tert-butyl (3R,4R)-3-(isopropylsulfonylamino)-4-[4-(trifluoromethylsulfonyloxy) phenyl] piperidine-1-carboxylate (8.5 g, 16.0 mmol, 1.00 eq) was dissolved in dioxane (200 mL) followed by the addition of potassium acetate (6.29 g, 64.1 mmol, 4.00 eq) and bis(pinacolato)diboron (12.2 g, 48.1 mmol, 3.00 eq). The reaction mixture was sparged with argon, followed by the addition of Pd(dppf)Cl_2_.DCM (1.96 g, 2.40 mmol, 0.150 eq). The reaction was stirred for 16 h at 90 ºC. After completion (based on TLC monitoring), the reaction mixture was evaporated and crude was purified by silica chromatography to afford tert-butyl (3R,4R)-3-(isopropylsulfonylamino)-4-[4-(4,4,5,5-tetramethyl-1,3,2-dioxaborolan-2-yl) phenyl] piperidine-1-carboxylate (7.5 g, 92% yield).

ESI m/z calculated: 508.3, Observed 509.1 [MH+]

**Step i: Synthesis of tert-butyl (3R,4R)-4-[4-(2-cyano-3-hydroxy-phenyl) phenyl]-3-(isopropyl sulfonylamino) piperidine-1-carboxylate**

A stirred suspension of tert-butyl (3R,4R)-3-(isopropylsulfonylamino)-4-[4-(4,4,5,5-tetramethyl-1,3,2-dioxaborolan-2-yl)phenyl] piperidine-1-carboxylate (7 g, 13.8 mmol, 1.00 eq), 2-bromo-6-hydroxy-benzonitrile (2.72 g, 13.8 mmol, 1.00 eq) and K_3_PO_4_ (8.76 g, 41.3 mmol, 3.00 eq) in dioxane (130 mL) and water (26 mL) was sparged with argon for 15 minutes followed by the addition of  Pd(PPh_3_)_4_ (1.59 g, 1.38 mmol, 0.100 eq) and the reaction mixture was heated at 90ºC for 16 h. After completion (monitored by TLC, R_f_=0.4, mobile phase: 70% ethyl acetate-hexane), the reaction mixture was diluted with ethyl acetate and washed with water, brine and dried over Na_2_SO_4_ and concentrated and purified by silica chromatography to afford tert-butyl (3R,4R)-4-[4-(2-cyano-3-hydroxy-phenyl) phenyl]-3-(isopropylsulfonylamino) piperidine-1-carboxylate, (3.6 g, 52% yield).

^1^H NMR (400 MHz, DMSO-*d_6_*): δ 11.17 (S,1H) 7.53-7.46 (m, 5H), 7.15 (d, J = 8.8Hz,1H), 7.02 (d, J = 8.4Hz,1H),6.92 (d, J = 8.4Hz,1H), 4.34 (s, 1H), 4.00 (d, 1H), 3.27 (m, 1H), 2.68 (m, 3H), 2.49 (m,1H), 1.73 (m, 2H), 1.41 (s, 9H ), 1.33 (m,1H), 0.93 (d, J = 6.8 Hz, 3H), 0.48 (d, J = 6.6 Hz, 3H) ppm­­­.

ESI m/z calculated 499.2, Observed 500.4 [MH+]

**Step j: Synthesis of N-[(3R,4R)-4-[4-(2-cyano-3-hydroxy-phenyl) phenyl]-3-piperidyl] propane-2-sulfonamide (BRD3290).**

To the stirred solution of tert-butyl (3R,4R)-4-[4-(2-cyano-3-hydroxy-phenyl) phenyl]-3-(isopropyl sulfonylamino) piperidine-1-carboxylate (9 g, 18.0 mmol, 1.00 eq) in dioxane (50 mL)  was added HCl (4 M in Dioxane, 45 mL, 180 mmol, 10.0 eq) and the reaction mixture was stirred at room temperature for 4 h. The reaction mixture was evaporated under reduced pressure and the residue triturated with ether to afford N-[(3R,4R)-4-[4-(2-cyano-3-hydroxy-phenyl)phenyl]-3-piperidyl]propane-2-sulfonamide, (6.5 g, 83% yield) as a white solid (hydrochloride salt).

^1^H NMR (400 MHz, DMSO-D6) : δ 11.27 (s,1H) ,9.12 (s,1H), 7.54-7.50 (m, 3H), 7.44-7.39 (m, 3H), 7.07 (d, J = 8 Hz, 1H), 3.74 (m, 1H), 3.47 (dd, J = 8.4 Hz, 1H), 3.34 (d, J = 8.4 Hz, 1H), 2.98 (t, J = 8 Hz, 1H), 2.84 (m, 2H), 2.34 (m, 1H), 2.11 (m, 1H), 1.98 (d, 1H), 0.93 (d, J = 6.8 Hz, 3H), 0.54 (d, J = 6.6 Hz, 3H) ppm.

^13^C NMR (151 MHz, DMSO) δ 161.30, 145.50, 142.27, 137.21, 134.20, 128.80, 128.11, 120.18, 116.30, 114.75, 98.05, 66.35, 52.40, 49.00, 45.08, 42.96, 29.11, 15.83, 15.56.

HRMS (ESI/QTOF) m/z: [M+H]^+^ Calcd for C_21_H_26_N_3_O_3_S^+^ 400.1689; found 400.1691.

Optical rotation: 0.238 ° (cell temperature: 25.00 °C)

Specific rotation (calc.): 45.820 °

**Synthesis of tetrahydropyran analogues**

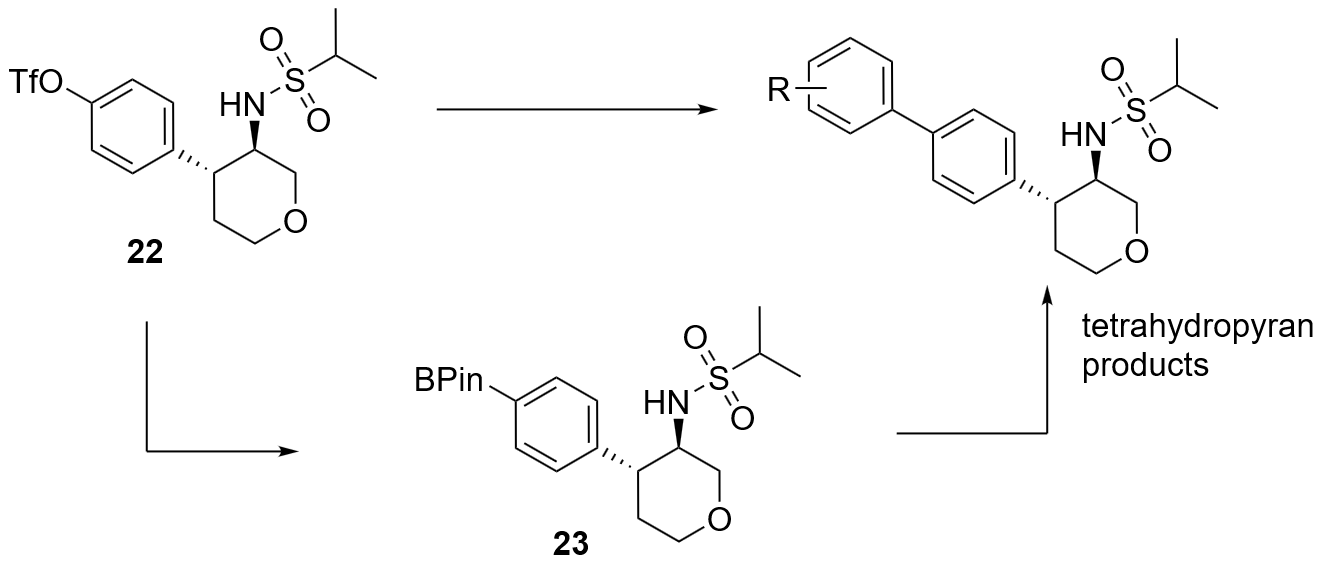

Compound 22 was prepared according to patent WO2008120093A1.

**Synthesis of N-((3R,4R)-4-(4-(4,4,5,5-tetramethyl-1,3,2-dioxaborolan-2-yl)phenyl)tetrahydro-2H-pyran-3-yl)propane-2-sulfonamide (23):**

A stirred solution of [4-[(3R,4R)-3-(isopropylsulfonylamino)tetrahydropyran-4-yl]phenyl] trifluoromethanesulfonate (900 mg, 2.09 mmol, 1.00 eq) in dioxane (10 mL) was sparged with argon for 10 minutes followed by the addition of bis(pinacolato) diboron (1.6 g, 6.26 mmol, 3.00 eq) and dry potassium acetate (1 g, 10.4 mmol, 5.00 eq). [1,1'-Bis(diphenylphosphino)ferrocene]dichloropalladium(II) (170 mg, 0.209 mmol, 0.100 eq) was added and the reaction mixture was heated with stirring at 100 ºC for 16 h. After completion of the reaction (monitored by TLC, 0.5 Rf in 50% ethyl aceate-hexane), the reaction mixture was filtered over celite. The filtrate was washed with water and brine and the organics were dried over anhydrous sodium sulfate and evaporated under reduced pressure to obtain the crude compound which was purified by flash chromatography over silica using 0-100% ethyl acetate-hexane to afford the target compound **23** (800 mg, 1.95 mmol, 94% yield). Boronates were not thoroughly characterized, but rather were used immediately in the subsequent cross-coupling step.

**Synthesis of tetrahydropyrans**

Tetrahydropyran products were prepared by coupling **22** or **23** with an aryl boronate or aryl halide or triflate respectively. In a typical procedure, a solution of the aryl triflate (1 eq) in dioxane (0.1 M) was sparged with argon folllowed by the addition of an aryl boronic acid (1.5 eq), potassium phosphate (2.5 eq) and tetrakis(triphenylphosphine)palladium(0) (10 mol%) and the reaction mixture was heated at 90 C until the conversion was judged complete by TLC. Filtration and evaporation of the reaction mixture afforded a crude residue that was first purified by flash chromatography over silica (hexanes/ethyl acetate) followed by preparative HPLC to furnish the pure products.

**Synthesis of piperidine compounds**

All piperidine compounds were prepared using analogous procedures to BRD3290, by Suzuki coupling between the appropriate aryl halide or triflate and boronic acid or ester, followed by purification with flash chromatography over silica (hexanes/EA) and resolution of the Boc-protected intermediates by chiral HPLC if required. Deprotection by treatment with HCl in dioxane generally afforded the products as hydrochloride salts. Data for the eutomers has been reported.

**Compound 1**

From (5-cyano-2-thienyl)boronic acid and **20**. Obtained as the hydrochloride.

^1^H NMR (400 MHz, DMSO-D6) : δ 8.91 (m, 2H), 8.0 (d, J = 3.92 Hz, 1H), 7.76 (d, J = 8.04 Hz, 2H), 7.68 (d, J = 3.88 Hz, 1H), 7.42-7.37 (m, 3H), 3.69-3.67 (m, 1H), 3.50-3.48 (m, 1H), 2.97 (m, 2H), 2.82-2.77 (m, 2H), 2.02-1.92 (m, 2H)0.94 (d, J = 6.72 Hz, 3H), 0.57 (d, J = 6.56 Hz, 3H).

LCMS: m/z = 390.33 [M+H]

**Compound 2**

From 4-hydroxyphenylboronic acid and **20**. Obtained as the hydrochloride.

^1^H NMR (DMSO-d6): δ 9.53 (s, 1H), 8.89 (brs, 1H), 7.55 (d, J = 8.1 Hz, 2H), 7.45 (d, J = 8.5 Hz, 2H), 7.37 (d, J = 9.0 Hz, 1H), 7.30 (d, J = 8.1 Hz, 2H), 6.85 (d, J = 8.6 Hz, 2H), 3.65-3.64 (m, 1H), 3.48-3.44 (m, 1H), 3.39-3.81 (m,1H), 2.94 (m, 1H), 2.80-2.74 (m, 2H), 2.38-2.32 (m, 1H), 2.07 (m, 1H), 1.95 (m, 1H), 0.92 (d, J = 6.8 Hz, 3H), 0.53 (d, J = 6.6 Hz, 3H).

LCMS m/z = 375.35 [M+H]

**Compound 3**

From **20** and 4-methoxyphenyl boronic acid. Obtained as the hydrochloride.

^1^H NMR (DMSO-d6): δ 9.01 (brs, 2H), 7.60-7.55 (m, 4H), 7.38-7.31 (m, 3H), 7.03 (d, J = 8.7 Hz, 2H), 3.79 (s, 3H), 3.69-3.66 (m, 1H), 3.48-3.44 (m, 1H), 3.34 (m, 1H), 2.94 (m, 1H), 2.80-2.73 (m, 2H), 2.39-3.35 (m, 1H), 2.07-2.03 (m, 1H), 1.95-1.92 (m, 1H), 0.92 (d, J = 6.8 Hz, 3H), 0.53 (d, J = 6.6 Hz, 3H).

LCMS m/z: 389.30 [M+H]

**Compound 4**

From (4-cyanophenyl)boronic acid and **20**. Obtained as the hydrochloride.

^1^H NMR (400 MHz, DMSO-d6): δ 8.92 (brs, 2H), 7.93 (d, J = 8.28 Hz, 2H), 7.86 (d, J = 8.36 Hz, 2H), 7.75 (d, J = 8.12 Hz, 2H), 7.42-7.38 (m, 3H), 3.72-3.68 (m, 1H), 3.51-3.47 (m, 1H), 2.99-2.94 (m, 1H), 2.84-2.76 (m, 2H), 2.45-2.40 (m, 1H), 2.07-1.94 (m, 2H), 0.93 (d, J = 6.76 Hz, 3H), 0.53 (d, J = 6.68 Hz, 3H).

LCMS: m/z = 384.2 [M+H]

**Compound 5**

From **21** and 5-bromo-2-oxo-1,2-dihydropyridine-4-carbonitrile. Obtained as the hydrochloride.

^1^H NMR (DMSO-d6): δ 7.64 (s, 1H), 7.43 -7-37 (m,4H) ,7.07 (s,1H), 6.91 (d,J=9.2 Hz, 1H) , 3.25 - 3.19 (m , 2H), 2.90 (m, 1H), 2.53 (m, 1H), 2.37-2.32 (m, 3H), 1.68 (s , 2H) , 0.90 (d, J=6-8Hz, 3H) , 0.49 (d, J=6.4Hz, 3H).

LCMS m/z: = 401.2 [M+H]

**Compound 6**

From 5-hydroxy-2-(4,4,5,5-tetramethyl-1,3,2-dioxaborolan-2-yl)benzonitrile and **20**. Obtained as the hydrochloride.

^1^H NMR (400 MHz, DMSO-d6): δ 10.35 (s, 1H), 8.92 (s, 2H), 7.50 (d, J = 8.0 Hz, 2H), 7.42 - 7.37 (m, 4H), 7.23 (d, J = 2.24 Hz, 1H), 7.18 (d, J = 8.48 Hz, 1H), 3.77 - 3.65 (m, 1H), 3.50 - 3.47 (m, 1H), 3.34 (s, 1H), 2.99 - 2.80 (m, 1H), 2.89 - 2.75 (m, 2H), 2.39 - 2.29 (m, 1H), 2.05 - 1.95 (m, 2H), 0.92 (d, J = 6.76 Hz, 3H), 0.48 (d, J = 6.64 Hz, 3H).

LCMS m/z = 400.30 [M+H]

**Compound 7**

**22** (90 mg, 0.209 mmol, 1.00 eq), 5-cyanothiophene-2-boronic acid (48 mg, 0.313 mmol, 1.50 eq) and Potassium Fluoride (61 mg, 1.04 mmol, 5.00 eq) were taken in a sealed tube and dissolved in MeOH:Toluene(1:1) (5 mL). The resulting mixture was sparged with argon for 30 min.

Palladium(II) acetate (2.8 mg, 0.01 mmol, 5 mol%) and XPHOS (10 mg, 0.02 mmol, 10 mol%) were added and the reaction was heated for 12 h at 95 ºC . After completion of the reaction (based on TLC monitoring) the reaction mixture was filtered using a sintered funnel and the filtrate was evaporated. The crude material was purified by flash chromatography.

^1^H NMR (400 MHz, DMSO-d6) δ 7.98 (d, J = 4.0 Hz, 1H), 7.71 – 7.66 (m, 3H), 7.48 (d, J = 8.0 Hz, 2H), 7.08 (d, J = 9.2 Hz, 1H), 4.02 –3.88 (m, 2H), 3.39 – 3.08 (m, 3H), 2.70 – 2.50 (m, 2H), 1.86 – 1.71 (m, 2H), 0.94 (d, J = 6.8 Hz, 3H), 0.50 (d, J = 6.8 Hz, 3H).

LCMS m/z-391.01 [M+H]

**Compound 8**

From **22** and 4-hydroxyphenyl boronic acid.

^1^H NMR (400 MHz, DMSO-d6): δ 9.49 (s, 1H), 7.49 (d, J = 7.96 Hz, 2H), 7.43 (d, J = 8.36 Hz, 2H), 7.37 (d, J = 8.08 Hz, 2H), 7.05 (d, J = 9.04 Hz, 1H), 6.83 (d, J = 8.36 Hz, 2H), 4.01-3.97 (m, 1H), 3.90-3.88 (m, 1H), 3.37 (m, 1H), 3.10 (t, J = 10.6 Hz, 1H), 2.66-2.63 (m, 1H), 2.44-2.42 (m, 2H), 1.89-1.83 (m, 1H), 1.75-1.72 (m, 1H), 0.92 (d, J = 6.76 Hz, 3H), 0.47 (d, J = 6.56 Hz, 3H).

LCMS m/z = 376.27 [M+H]

**Compound 9**

From 4-methoxyphenylboronic acid and **22**.

^1^H NMR (400 MHz, DMSO-d6): δ 7.55 (t, J = 8.8 Hz, 4H), 7.40 (d, J = 8.08 Hz, 2H), 7.01 (d, J = 8.52 Hz, 2H), 4.01-3.98 (m, 1H), 3.91-3.89 (m, 1H), 3.79 (s, 3H), 3.31 (m, 1H), 3.11(t, J = 10.3 Hz, 1H), 2.69-2.64 (m, 1H), 2.45-2.43 (m, 2H), 1.90-1.84 (m, 1H), 1.76-1.73 (m, 1H), 0.90 (d, J = 6.84 Hz, 3H), 0.47 (d, J = 6.56 Hz, 3H).

LCMS m/z = 390.18 [M+H]

**Compound 10**

From (4-cyanophenyl)boronic acid and **22**.

^1^H NMR (400 MHz, DMSO-d6): δ 7.92-7.85 (m, 4H), 7.69 (d, J = 8.4 Hz, 2H), 7.5 (d, J = 8 Hz, 2H), 7.08 (s, 1H), 4.03-3.99 (m, 1H), 3.92-3.89 (m , 1H), 3.38-3.36 (m, 2H), 3.12 (t, J = 10.6 Hz, 1H), 2.75-2.68 (m, 1H), 1.93-1.89 (m, 1H), 1.77-1.74 (m, 1H), 0.92 (d, J = 6.8 Hz, 3H), 0.47 (d, J = 6.4 Hz, 3H).

LCMS m/z = 385.2 [M+H]

**Compound 11**

From **23** and 5-bromo-2-oxo-1,2-dihydropyridine-4-carbonitrile.

^1^H (400 MHz, DMSO-d6): δ 7.60 (brs, 1H), 7.44 (m, 4H), 7.0 (m, 1H), 6.69-6.67 (m, 1H), 4.0-3.97 (m, 1H), 3.91-3.87 (m, 1H), 3.37-3.34 (m, 2H), 3.13-3.08 (m, 1H), 2.73-2.66 (m, 1H), 2.40-2.32 (m, 1H), 1.94-1.85 (m, 1H), 1.75-1.72 (m, 1H), 0.92 (d, J=6.8 Hz, 3H), 0.49 (d, J=6.4 Hz, 3H).

LCMS (m/z) = 402.2 [M+H]

**Compound 12**

From 5-hydroxy-2-(4,4,5,5-tetramethyl-1,3,2-dioxaborolan-2-yl)benzonitrile and **22**.

^1^H NMR (400 MHz, DMSO-d6): δ 7.49-7.43 (m, 4H), 7.38 (d, J = 8.52 Hz, 1H), 7.20-7.19 (m, 1H), 7.17-7.15 (m, 2H), 4.01-3.97 (m, 1H), 3.92-3.89 (m, 1H), 3.38-3.35 (m, 2H), 3.11 (t, J = 10.6 Hz, 1H), 2.71-2.67 (m, 1H), 2.41-2.38 (m, 1H), 1.90 (m, 1H), 1.78 (m, 1H), 0.92 (d, J = 6.8 Hz, 3H), 0.48 (d, J = 6.6 Hz, 3H).

LCMS m/z = 401.34 [M+H]

**Compound 13**

From 4-bromo-1,3-benzoxazole and **21**. After cross-coupling, the boc-protected intermediate was deprotected via the following procedure: tert-butyl (3R,4R)-4-[4-(1,3-benzoxazol-4-yl)phenyl]-3-(isopropylsulfonylamino)piperidine-1-carboxylate (25 mg, 0.05 mmol, 1.00 eq) was dissolved in dichloromethane (2 mL) under an atmosphere of nitrogen and the solution was cooled to 0ºC with stirring. Trimethylsilyl trifluoromethanesulfonate (0.04 mL, 0.200 mmol, 4.00 eq) was added, followed by triethylamine (0.03 mL, 0.25 mmol, 5.00 eq) and the reaction was stirred for 2h at 0ºC. The reaction was quenched by the addition of water, the organics were diluted with 2 mL additional dichloromethane, separated, dried over sodium sulfate and evaporated to afford a crude residue which was purified by preparative HPLC. Obtained as the acetate salt.

^1^H NMR (400 MHz, DMSO-d6): δ 8.79 (s, 1H), 7.92 (d, J = 8.08 Hz, 2H), 7.74 (d, J = 8.0 Hz, 2H), 7.61 (d, J = 7.68 Hz, 1H), 7.51 (t, J = 7.88 Hz, 2H), 7.43 (t, J = 8.88 Hz, 1H), 3.25 - 3.16 (m, 1H), 2.94 (d, J = 12.04 Hz, 1H), 2.61 -2.53 (m, 1H), 2.41 - 2.24 (m, 1H), 1.74 (d, J = 5.48 Hz, 2H), 0.92 (d, J = 6.8 Hz, 3H), 0.50 (d, J = 8.08 Hz, 3H).

LCMS m/z = 400.00 [M+H]

**Compound 14**

From 2-bromo-3-fluoro-6-hydroxy-benzonitrile and **21**. After deprotection, the compound was further purified by preparative HPLC (acetonitrile/water + 0.05% TFA), which afforded the compound as the trifluoroacetate.

^1^H NMR (400MHZ,DMSO) : δ 7.44-7.39 (m, 5H), 7.00-6.95 (m, 2H), 3.24-3.21 (m, 2H), 2.97 (s, 1H), 2.66-2.59 (m, 1H), 2.44-2.39 (m, 1H), 2.32-2.24 (m, 1H), 1.75-1.74 (m, 2H), 0.89 (d, J=6.8, 3H), 0.53 (d, J=6.8, 3H).

LCMS m/z = 418.26 [M+H]

**Compound 15**

From 2-bromo-4-fluoro-6-hydroxy-benzonitrile and **21**. Obtained as the hydrochloride.

^1^H NMR (400 MHz, DMSO-d6): δ 11.83 (s, 1H), 8.91 (s, 1H), 7.58-7.56 (d, J = 8 Hz, 2H), 7.44-7.39 (m, 3H), 6.84-6.82 (d, J = 9.2 Hz ,2H), 3.69-3.68 (m, 1H), 3.48-3.46 (m, 1H),3.36 (s,1H), 2.99-2.94 (m,1H), 2.81-2.79 (m, 2H), 2.35-2.31 (m,1H), 2.06-2.00 (m, 2H), 0.93-0.91 (d, J = 6.8 Hz, 3H), 0.55-0.53 (d, J = 6.8 Hz, 3H) ppm.

LCMS m/z = 418.2 [M+H]

**Compound 16**

From 6-bromo-3-fluoro-2-hydroxy-benzonitrile and **21**. Obtained as the hydrochloride.

^1^H NMR (400 MHz, DMSO-d6): δ 11.37 (s, 1H), 8.8 (s, 1H), 7.88-7.85 (d, J = 11.6 Hz, 1H), 7.52-7.50 (d, J = 7.6 Hz, 2H), 7.43-7.39 (t, J = 16 Hz ,3H), 7.06-7.04 (d, J = 8.4 Hz, 1H), 3.71-3.65 (m, 1H), 3.48-3.46 (m, 1H),3.36 (s,1H), 2.99-2.94 (m,1H), 2.81-2.79 (m, 2H), 2.35-2.31 (m,1H), 2.06-2.00 (m, 2H), 0.93-0.91 (d, J = 6.8 Hz, 3H), 0.55-0.53 (d, J = 6.8 Hz, 3H) ppm.

LCMS m/z = 418.3 [M+H]

**Compound 24**

From 2-bromo-4,6-dimethoxy-benzonitrile and **21**. Obtained as the trifluoroacetate after preparative HPLC.

^1^H NMR (400MHZ,DMSO) : δ 7.49 (d, J=7.6Hz, 2H), 7.41 (d, J=8, 2H), 6.91 (d, J=9.2 Hz, 1H), 6.75 (s, 1H), 6.60 (s, 1H), 3.93 (s, 3H), 3.89 (s, 3H), 3.25-3.19 (m, 2H), 2.92 (d, J=12Hz, 1H), 2.66-2.53 ( m, 1H), 2.38-2.27 ( m, 3H), 1.70 ( Brs, 2H), 0.90 (d, J=6.8Hz, 3H), 0.51 (d, J=6.4Hz, 3H) .

LCMS m/z = 444.3 [M+H]

**Compound 25**

From 6-bromo-2,3-dimethoxy-benzonitrile and **21**. Obtained as the trifluoroacetate after preparative HPLC.

^1^H NMR (400MHZ,DMSO) : δ 7.47-7.44 (m, 5H), 7.22 (d, J=8.8 Hz, 1H), 6.92 (d, J=9.2 Hz, 1H), 3.91 (d, J=12.4 Hz, 6H), 3.32-3.18 (m, 2H), 2.92-2.89 (m, 1H), 2.56-2.28 (m, 4H), 1.71-1.70 (m, 2H), 0.90 (d, J=6.8, 3H), 0.49 ( d, J=6.4, 6H).

LCMS m/z = 444.2 [M+H]

**Compound 26**

From 6-bromo-2-hydroxy-3-methoxy-benzonitrile and **21**. Obtained as the hydrochloride.

^1^H-NMR (400 MHz, DMSO-d6): δ 10.54 (br s, 1H), 8.99 (br s, 1H), 7.59 (d, J=8.00 Hz, 2H), 7.43-7.36 (m, 3H), 7.31 (d, J=8.80 Hz, 1H), 6.89 (d, J=8.40 Hz, 1H), 3.89 (s, 3H), 3.72-3.66 (m, 1H), 3.46 (dd, J=4.2, 12.2 Hz, 1H), 3.35-3.32 (m, 1H), 2.99-2.93 (m, 1H), 2.83-2.75 (m, 2H), 2.34-2.27 (m, 1H), 2.07-1.95 (m, 2H), 0.92 (d, J=6.80 Hz, 3H), 0.54 (d, J=6.80 Hz, 3H).

LCMS m/z = 430.2 [M+H]

**Compound 27**

From 2-bromo-6-hydroxy-4-methoxy-benzonitrile and **21**. Obtained as the hydrochloride.

^1^H-NMR (400 MHz, DMSO-d6): δ 11.22 (br s, 1H), 8.95-8.91 (m, 1H), 7.54 (d, J=8.12 Hz, 2H), 7.43-7.37 (m, 3H), 6.56-6.55 (m, 1H), 6.47-6.46 (m, 1H), 3.81 (s, 3H), 3.71-3.65 (m, 1H), 3.49-3.46 (m, 1H), 3.36-3.33 (m, 1H), 2.99-2.93 (m, 1H), 2.83-2.77 (m, 2H), 2.32-2.28 (m, 1H), 2.06-1.94 (m, 2H), 0.92 (d, J=6.80 Hz, 3H), 0.54 (d, J=6.80 Hz, 3H).

LCMS m/z = 430.2 [M+H]

**Compound 28**

From 6-bromo-2-hydroxy-3-methyl-benzonitrile and **21**. Obtained as the hydrochloride.

^1^H NMR (400MHZ,DMSO) : δ 10.19 (Brs, 1H), 8.94 (Brs, 1H), 7.52-7.50 (m, 2H), 7.50-7.37 (m, 4H), 6.88 (d, J=7.6, 5H), 3.70 (Brs, 1H), 3.46 (d, J=8.8, 1H), 2.96 (t, J=10.8 Hz, 1H), 2.80-2.78 (m, 1H), 2.32-2.26 ( m, 4H), 2.07-1.98 ( m, 3H), 0.92 ( d, J=6.8, 3H), 0.53 ( d, J=5.6, 3H).

LCMS m/z = 414.3 [M+H]

**Compound 29**

From 6-bromo-3-hydroxy-2-methoxy-benzonitrile and **21**. Obtained as the trifluoroacetate after preparative HPLC.

^1^H NMR (400 MHz, DMSO-D6) : δ 8.97 (s, 2H), 7.52 (d, J = 8 Hz, 2H), 7.42 (m, 4H), 7.31 (s, 1H), 4.67 (t, J = 8.8 Hz, 2H), 3.70 (m, 1H), 3.48 (m, 1H), 2.98 (m, 1H), 2.82 (m, 2H), 2.34 (m, 1H), 2.08 (m, 2H), 0.93 (d, J = 6.8 Hz, 3H), 0.55 (d, J = 6.6 Hz, 3H).

LCMS: m/z = 426.3 [M+H]
