## Supplemental Tables for "Discovery of GluA3 preferring AMPA receptor positive allosteric modulator BRD3290"

| Compound | 1 | 2 | 3 | 4 | 5 | 6 | 7 | 8 | 9 | 10 | 11 | 12 | 13 | 14 | 15 | 16 | BRD3290 | 24 | 25 | 26 | 27 | 28 | 29 |
| --- | --- | --- | --- | --- | --- | --- | --- | --- | --- | --- | --- | --- | --- | --- | --- | --- | --- | --- | --- | --- | --- | --- | --- |
| GluA3i/3i | ND | -0.52 | -0.02 | ND | -1.03 | 1.02 | 0.65 | 0.53 | 0.25 | ND | 0.68 | 1.49 | -0.15 | -0.91 | -1.20 | -0.78 | 0.64 | 0.43 | 0.41 | -0.58 | ND | 0.42 | ND |
| GluA3i/2o | -0.33 | -0.50 | -0.21 | -0.84 | -0.52 | 1.19 | 0.86 | 1.00 | -0.21 | -0.13 | 0.60 | 2.09 | 0.18 | -0.16 | -0.52 | -0.01 | 0.72 | 0.35 | 0.48 | -0.14 | -0.35 | 0.45 | -0.34 |
| GluA3i/2i | -0.64 | -0.67 | -0.46 | -0.63 | -0.94 | 0.70 | 0.68 | 0.82 | -0.28 | 0.21 | 0.31 | 1.66 | -0.44 | -0.78 | -0.99 | -0.56 | -0.20 | 0.02 | 0.18 | -0.49 | -1.18 | 0.04 | -1.03 |
| GluA3o/2i | ND | -0.95 | -0.33 | ND | -0.93 | 0.85 | 0.58 | -0.13 | -0.13 | ND | 0.12 | 1.72 | 0.08 | -0.37 | -0.72 | 0.06 | 0.46 | 0.47 | 0.67 | -0.17 | ND | 0.62 | ND |
| GluA3o/2o | -0.67 | -0.74 | -0.42 | -1.11 | -0.79 | 1.16 | 0.63 | 0.57 | -0.23 | -0.19 | -0.06 | 2.13 | 0.10 | -0.12 | -0.63 | 0.45 | 0.45 | 0.43 | 0.78 | -0.25 | -0.11 | 0.32 | -0.72 |
| GluA2o/2o | ND | -0.93 | -0.52 | ND | -0.86 | 1.49 | 1.08 | -0.35 | -0.64 | ND | 0.39 | 1.59 | 0.40 | 0.22 | -0.37 | 0.47 | 1.20 | 0.49 | 0.68 | -0.28 | ND | 0.60 | ND |
| GluA1i/2o | -0.43 | -0.67 | -0.38 | -0.58 | -0.91 | 0.70 | 0.77 | 0.78 | -0.21 | 0.29 | -0.02 | 1.79 | -0.28 | -0.66 | -0.83 | -0.12 | -0.11 | 0.50 | 0.64 | -0.46 | -0.60 | 0.32 | -0.39 |
| GluA1i/1i | ND | ND | ND | ND | -0.96 | 0.58 | ND | ND | ND | ND | 0.06 | 1.73 | -0.12 | ND | ND | ND | 0.20 | ND | ND | ND | ND | ND | ND |

**Supplemental Table S1:** Single delta-log measurements for compound activity compared to HBT-1. ND = not determined.

| Compound | 1 | 2 | 3 | 4 | 5 | 6 | 7 | 8 | 9 | 10 | 11 | 12 | 13 | 14 | 15 | 16 | BRD3290 | 24 | 25 | 26 | 27 | 28 | 29 |
| --- | --- | --- | --- | --- | --- | --- | --- | --- | --- | --- | --- | --- | --- | --- | --- | --- | --- | --- | --- | --- | --- | --- | --- |
| GluA3i/3i | ND | 0.14 | 0.36 | ND | -0.11 | 0.32 | -0.13 | -0.25 | 0.46 | ND | 0.70 | -0.30 | 0.13 | -0.25 | -0.37 | -0.66 | 0.75 | -0.08 | -0.22 | -0.12 | ND | 0.09 | ND |
| GluA3i/2o | 0.10 | 0.16 | 0.17 | -0.26 | 0.40 | 0.50 | 0.09 | 0.23 | 0.01 | -0.42 | 0.62 | 0.30 | 0.46 | 0.50 | 0.31 | 0.10 | 0.83 | -0.15 | -0.15 | 0.31 | 0.25 | 0.12 | 0.05 |
| GluA3i/2i | -0.21 | 0.00 | -0.09 | -0.05 | -0.03 | 0.00 | -0.09 | 0.05 | -0.07 | -0.08 | 0.32 | -0.13 | -0.15 | -0.12 | -0.16 | -0.44 | -0.09 | -0.49 | -0.45 | -0.03 | -0.58 | -0.29 | -0.64 |
| GluA3o/2i | ND | -0.28 | 0.04 | ND | -0.01 | 0.15 | -0.19 | -0.90 | 0.08 | ND | 0.14 | -0.07 | 0.36 | 0.29 | 0.11 | 0.17 | 0.57 | -0.04 | 0.04 | 0.29 | ND | 0.30 | ND |
| GluA3o/2o | -0.24 | -0.07 | -0.05 | -0.53 | 0.12 | 0.47 | -0.14 | -0.21 | -0.02 | -0.48 | -0.05 | 0.34 | 0.38 | 0.53 | 0.21 | 0.57 | 0.56 | -0.08 | 0.15 | 0.21 | 0.49 | 0.00 | -0.33 |
| GluA2o/2o | ND | -0.26 | -0.15 | ND | 0.05 | 0.79 | 0.30 | -1.13 | -0.43 | ND | 0.41 | -0.20 | 0.68 | 0.88 | 0.46 | 0.59 | 1.30 | -0.02 | 0.05 | 0.18 | ND | 0.28 | ND |
| GluA1i/2o | 0.00 | 0.00 | 0.00 | 0.00 | 0.00 | 0.00 | 0.00 | 0.00 | 0.00 | 0.00 | 0.00 | 0.00 | 0.00 | 0.00 | 0.00 | 0.00 | 0.00 | 0.00 | 0.00 | 0.00 | 0.00 | 0.00 | 0.00 |
| GluA1i/1i | ND | ND | ND | ND | -0.05 | -0.12 | ND | ND | ND | ND | 0.08 | -0.06 | 0.17 | ND | ND | ND | 0.31 | ND | ND | ND | ND | ND | ND |

**Supplemental Table S2**: Double delta-log measurements for compound selectivity over GluA1i/2o. ND = not determined.

**Supplemental activity and selectivity data for compounds 24-29**

| R-group | 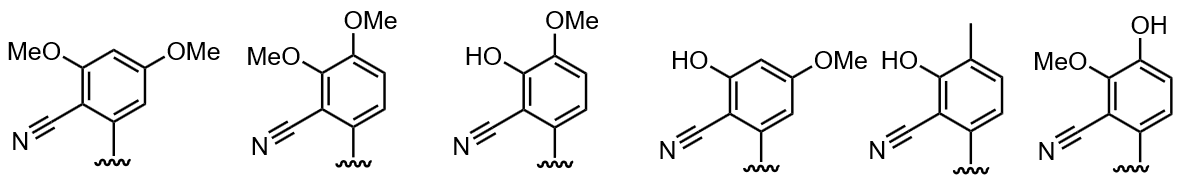 | | | | | |
| --- | --- | --- | --- | --- | --- | --- |
| Compound no. | **24** | **25** | **26** | **27** | **28** | **29** |
| Max selectivity | 0.9 x | 1 x | 2 x | 3 x | 2 x | 1 x |
| Average Δlog activity vs. HBT-1 | 0.38 | 0.55 | -0.34 | -0.56 | -0.40 | -0.62 |

**Supplemental Table S3:** Additional structure-activity relationship data for BRD3290 analogues. Max selectivity is the highest fold selectivity observed for any GluA3-containing heterodimer compared to GluA1i/2o. Average Δlog vs. HBT-1 is the arithmetic mean of all single Δlog unit activity measurements referenced to HBT-1 across all cell lines measured (a Δlog value of 1.0 indicates a compound that is 10 x more active than HBT-1).

**Supplemental Table 4:** In vitro ADME data for BRD3290

| **In vitro ADME Properties of BRD3290** | |
| --- | --- |
| Mouse microsome t_1/2_ / min | >145 |
| Mouse hepatocyte Cl_int_ (µL/min/million cells) | 67.7 |
| Permeability A-B (MDR1-MDCKII)/10^-6^cm s^-1^ | 0.36 |
| Efflux ratio | 4.70 |
| Kinetic solubility / µM  (PBS (pH 7.4) + 1% DMSO | 99.9 |
| Plasma t_1/2_ / hours (5mg/kg dosed IV) | 1.15 |
| Clearance / mL/min/kg  (5mg/kg dosed IV) | 11.72 |
